# A transfer RNA methyltransferase with an unusual domain composition catalyzes 2′-*O*-methylation at position 6 in tRNA

**DOI:** 10.1101/2025.04.03.647152

**Authors:** Teppei Matsuda, Ryota Yamagami, Aoi Ihara, Takeo Suzuki, Akira Hirata, Hiroyuki Hori

## Abstract

*Thermococcus kodakarensis* tRNA^Trp^ contains 2′-*O*-methylcytidine at position 6 (Cm6). However, the tRNA methyltransferase responsible for the modification has not been identified. Using comparative genomics we predicted TK1257 as a candidate for the modification. Biochemical and mass spectrometry studies of purified recombinant TK1257 gene product show it to possess a tRNA methyltransferase activity for Cm6 formation. This protein has a highly unusual composition of domains, containing N-terminal ferredoxin-like, SPOUT catalytic and THUMP domains. Previous to this study, all known THUMP-related tRNA methyltransferases were shown to contain a Rossmann fold catalytic domain and the nucleosides they produced were *N*2-methylguanosine and/or *N*2, *N*2-dimethylguanosine. Therefore, our findings extend the knowledge of architecture of tRNA methyltransferases. We named the TK1257 gene product TrmTS and showed it can synthesize Am6 and Um6 as well as Cm6. A *trmTS* gene deletion strain showed slight growth retardation at high temperatures. Site-directed mutagenesis studies based on structural model revealed catalytically and structurally important amino acid residues in TrmTS and identified a TrmTS-specific linker is structurally essential. We showed that TrmTS recognizes the 3′-CCA terminal region and a stretch loop connected with at least two stems in RNA. Finally, we constructed a model of the binding between TrmTS and tRNA.

## INTRODUCTION

To date, more than one hundred modified nucleosides have been discovered in tRNA (1–3). Of the modified nucleosides in tRNA, methylated nucleosides are the most abundant (4). Almost all methylated nucleosides in tRNA are synthesized by site-specific tRNA methyltransferases (4).

Although a few RNA methyltransferases use N5, N10-methylenetetrahydrofolate as a methyl group donor (5–7), the majority of RNA methyltransferases use S-adenosyl-L-methionine (SAM) (4). SAM-dependent methyltransferases can be classified into more than five classes by the structures of their catalytic domain (8). The majority of SAM-dependent tRNA methyltransferases are classified as the class I enzymes, which possess a Rossmann fold catalytic domain. The classes II, III and V enzymes methylate low molecular weight compounds or protein and do not methylate RNAs. The majority of class IV enzymes methylate RNA and possess the SpoU-TrmD (SPOUT) catalytic domain (9–13). SpoU is a classical name of TrmH (14, 15) and remains in the superfamily name. TrmD is the tRNA methyltransferase for synthesis of 1-methylguanosine at position 37 (m^1^G37) (16, 17). In addition, TrmO (a SAM-dependent tRNA methyltransferase for N6-methylation in N6-threonylcarbamoyladenosine synthesis) exceptionally possesses a β-barrel catalytic domain (18).

The existence of the SPOUT RNA methyltransferase superfamily (class IV enzymes) was initially predicted by a bioinformatic study (9) and then confirmed by structural studies (19–25). The SPOUT catalytic domain of Class IV enzymes contains a trefoil-knot structure, in which its C-terminal polypeptide threads through a polypeptide loop. Three conserved amino acid sequence motifs (motifs 1, 2 and 3) have been proposed in the catalytic domain of SPOUT RNA methyltransferase superfamily (9). In the cases of TrmH and a subset of other SPOUT RNA 2′-*O*-methyltransferases such as TrmL, the catalytic arginine residue is contained in the motif 1 and conserved amino acid residues in the motif 2 and motif 3 contribute to formation of the trefoil-knot structure and SAM-binding pocket (26–28).

The SPOUT RNA methyltransferases can be further classified by their products such as 2′-*O*-methylated nucleosides, m^1^G, 1-methyladenosine (m^1^A), and 1-methylpseudouridine (m^1^Ψ) (11). Within the SPOUT RNA 2′-*O*-methyltransferases, four tRNA 2′-*O*-methyltransferase families [TrmH (14, 15, 29), TrmL (30), TrmJ (31) and Trm56 (32, 33) families] have been identified. TrmH catalyzes the methylation of 2′-OH of ribose of G18 and produces 2′-*O*-methylguanosine at position 18 (Gm18) in tRNA (34–36). Likely, TrmL, TrmJ, and Trm56 catalyze the 2′-*O*-methylation at positions 34, 32, and 56, respectively, and produces 2′-*O*-methylcytidine (Cm34) and 5-carboxylmethylaminomethyl-2′-*O*-methyluridine (cmnm^5^Um34), Cm32 and 2′-*O*-methyluridine (Um32), and Cm56, respectively (30, 32, 33, 37, 38).

In general, 2′-*O*-methylation in RNA tilts the equilibrium of ribose puckering from the C2′-endo form to the C3′-endo form (39). Therefore, 2′-*O*-methylation stabilizes the local structure of RNA. Furthermore, because 2′-*O*-methylation prevents cleavage by RNases, it may prolong the half-life of RNA (40). In the case of thermophilic archaea, 2′-*O*-methylations at multiple positions in tRNA coordinately stabilize the L-shaped tRNA structure (41). In fact, thermophile-specific modified nucleosides often contain 2′-*O*-methylation, for example 1, 2′-*O*-dimethylinosine (m^1^Im) and *N*^2^, *N*^2^, 2′-*O*-trimethylguanosine (m^2^_2_Gm) (42). Furthermore, 2′-*O*-methylations in tRNAs are involved in higher biological phenomena such as stress-resistance or stress-response (43–46), immune response (47–53) and human diseases (54).

RNA modification enzymes often contain an RNA binding domain(s). The thiouridine synthetase, methyltransferase and pseudouridine synthase (THUMP) domain is a typical RNA binding domain (55) and has only been found in tRNA modification enzymes (56). THUMP-related tRNA modification enzymes can be classified into five types, namely 4-thiouridine synthase (57–59), deaminase (60), methyltransferase (61–66), a partner protein of acetyltransferase (67, 68) and pseudouridine synthase (69–72). Although the THUMP domain can fold autonomously, its affinity for tRNA is very low (73). The crystal structure of *Bacillus anthracis* TtuI (previous name ThiI (74); tRNA 4-thiouridine synthetase) revealed that the THUMP domain is composed of α-helices and β-strands (75) as predicted by a bioinformatics study (55). Furthermore, in this report (75), a tRNA-binding model was constructed and the THUMP domain of TtuI was placed near the CCA-terminus of substrate tRNA because CCA terminus is essential for the sulfur-transfer reaction by TtuI (76). THUMP domain recognizes the 3′-end of RNA (*i.e.,* CCA terminus of tRNA). This concept was established by numerous biochemical and structural studies of THUMP-related tRNA modification enzymes (60, 63, 75, 77–81). The N-terminal ferredoxin-like domain adjusts the angle and distance between the THUMP and catalytic domains (79). Furthermore, it should be mentioned that a THUMP-related structure is contained in an accessory domain of human PUS10 (tRNA pseudouridine synthase for Ψ54 and Ψ55 in tRNA) (82) and that the CCA terminus in tRNA is not essential for pseudouridine modification by PUS10 (83). All THUMP-related tRNA methyltransferases discovered are tRNA N2-guanine methyltransferases, which produce N2-methylguanosine (m^2^G) and/or N2, N2-dimethylguanosine (m^2^_2_G) (56).

In 2019, we reported the nucleotide sequence of tRNA^Trp^ from *Thermococcus kodakarensis*, a hyper-thermophilic archaeon (Figure 1A, ref. 84). This tRNA^Trp^ contains Cm6. However, the tRNA methyltransferase responsible for the synthesis of Cm6 has not been identified yet. In this study, we have found that a SPOUT 2′-*O*-methyltransferase with a THUMP domain produces the Cm6 modification in tRNA. This tRNA methyltransferase, which has a very unusual domain composition, acts on the 2′-*O*-methylation of aminoacyl-stem in tRNA. In this article, we report the identification of the gene encoding this tRNA methyltransferase, its enzymatic properties, and predicted structure as well as the presence of conserved amino acid residues in the catalytic domain and proposed tRNA-binding mode.

**Figure 1.**
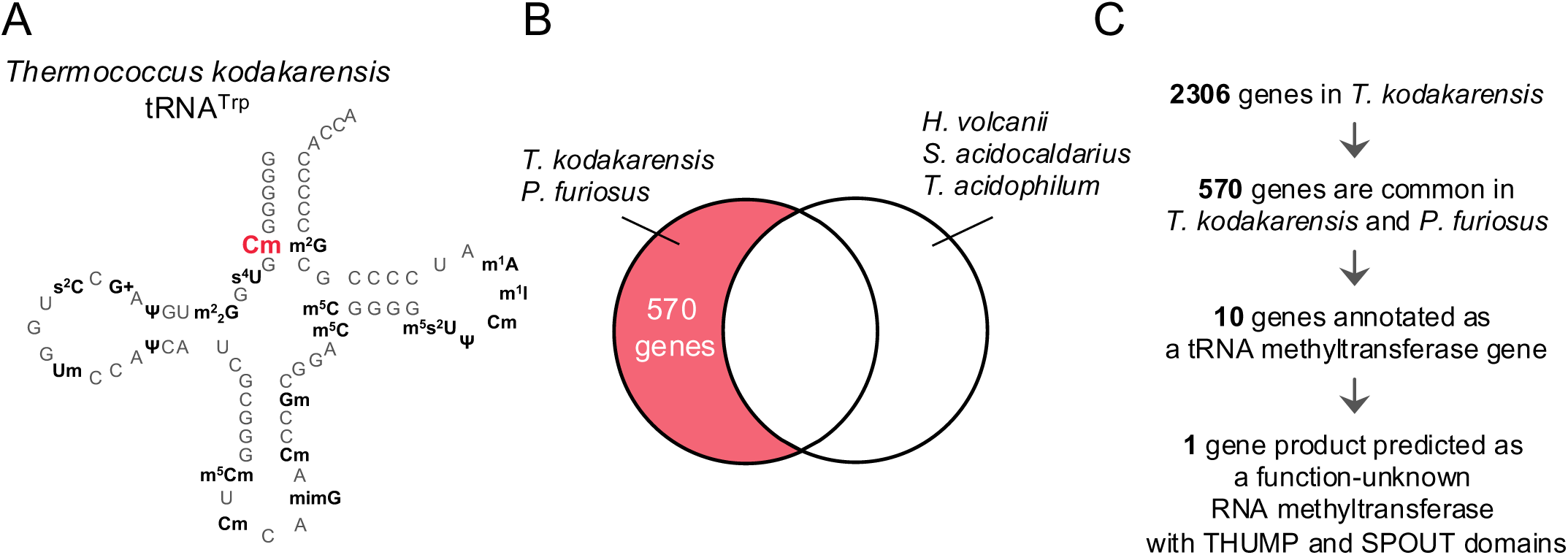
Nucleotide sequence of *Thermococcus kodakarensis* tRNA^Trp^ and scheme of comparative genomics. A, The secondary structure of *T. kodakarensis* tRNA^Trp^ is depicted as a cloverleaf structure. The Cm6 is highlighted in red. B, The responsible gene for the Cm6 modification in tRNA exists in *T. kodakarensis* and *P. furiosus* genomes was predicted by comparative genomics. *H. volcanii*, *S. acidocaldarius* and *T. acidophilum* do not possess the Cm6 modification in tRNA. The 570 genes were common in *T. kodakarensis* and *P. furiosus*. C, Scheme of comparative genomics is illustrated. An explanation of details is described in the main text.

## MATERIALS AND METHODS

### Materials

[Methyl-^3^H]-SAM (2.47 TBq/mmol) and [Methyl-^14^C]-SAM (1.95 GBq/mmol) were purchased from PerkinElmer. Non-radioisotope-labeled SAM was obtained from Sigma. DNA oligonucleotides were obtained from Thermo Fisher Scientific.

### Strains, media, and culture conditions

*Thermococcus kodakarensis* strains and plasmids are listed in Supplementary Table 1. *Thermococcus kodakarensis* KUW1 strain was a kind gift from Dr. Tamotsu Kanai (Toyama Prefectural University, Japan; ref. 85). *Thermococcus kodakarensis* KUWA strain, which was used for the construction of the TK1257 gene deletion (*ΔTK1257*) strain, is auxotrophic for agmatine (86). All *T. kodakarensis* strains were cultivated under anaerobic conditions at 85°C or 93°C in nutrient-rich medium (MA-YT). MA-YT medium (1 L) contains 0.8 × Marine Art SF1 reagent (Osaka Yakken Co. Ltd., Osaka, Japan), 5 g yeast extract (Y), and 5 g tryptophan (T). 2 g elemental sulfur (S^0^) or 5 g sodium pyruvate (Pyr) and 5 g Amycol #3-L (Mdx) was supplied into 1 L MA-YT medium prior to culture. Amycol #3-L (Nippon Starch Chemical, Osaka, Japan) contains a mixture of malto-oligosaccharides with lengths from 1 to 12 glucose units. Growth characteristics of the KUW1 and the *ΔTK1257* strains were examined in MA-YT-Pyr-Mdx medium. Growth at 85°C and 93 °C was examined by monitoring the optical density at 660 nm. For all liquid media, resazurine (final concentration, 0.5 mg/L) was supplemented as an oxygen indicator, and 5.0% Na_2_S H_2_O was added until the medium became colorless. For colony isolation, solid MA-YT media containing 1 g of gelrite and 0.4 g of polysulfide per 0.1 L were used. *Escherichia coli* DH5α strain was used for construction of the gene disruption plasmids and was grown at 37 °C in LB medium containing ampicillin (final concentration, 100 mg/L). Unless otherwise described in this study, chemicals were purchased from Nacalai Tesque (Kyoto, Japan) except for the gelrite, which was obtained from Wako Pure Chemicals (Osaka, Japan),

### Comparative genomic analysis

The genomes used in this study were obtained from the complete genomes available in the BioProject of the NCBI database. Protein sequence analyses were performed using BLASTP with an inclusion threshold *E*-value of 1e-10.

### Construction of TK1257 gene product expression vector in *Escherichia coli* cells

*Thermococcus kodakarensis* strain KUW1 genomic DNA was extracted from the cells with a phenol-chloroform mixture. TK1257 gene was amplified by polymerase chain reaction (PCR) from the genomic DNA using the following primers: TK1257_F, 5′-GAA GGA GAT ATA <u>CAT ATG</u> AAG TTT CTC GTC AAG ACT CAG AGG- 3′; TK1257_R, 5′-GAG CTC GAA TTC <u>GGA</u> <u>TCC</u> TTA TCA GGA AGA GTC CTC GGC TTT TTC- 3′. The underlined sequences show the restriction enzyme NdeI and BamHI cleavage sites, respectively. The amplified TK1257 gene was cloned into a pET30a vector using NEBuilder HiFi DNA Assembly Master Mix (New England Biolabs) in accordance with the manufacturer’s instruction.

### Expression of TK1257 gene product in *E. coli* cells

The TK1257 expression vector was introduced into *E. coli* BL21 (DE3) Rosetta 2 strain cells. The transformants were cultivated in 250 ml of LB liquid medium containing 50 µg/ml of kanamycin at 37°C for 12 h, then added into 1 L of the same medium, and the culture was continued at 37°C. When the optical density at 600 nm (OD_600 nm_) reached ∼0.8, isopropyl-ß-D-thiogalactopyranoside (IPTG) was added into the medium to a final concentration of 1 mM and then the culture was continued at 37°C for 4 h. The cells were collected by centrifugation at 10,000 x g at 4°C for 20 min, frozen by liquid nitrogen, and stored at -80°C.

### Purification of TK1257 gene product

0.7 g of wet cells were suspended in 10 mL buffer A [50 mM Tris-HCl (pH 7.6), 5 mM MgCl_2_, 200 mM KCl, 6 mM 2-mercaptoethanol and 5% (v/v) glycerol] supplemented with 100 μL protease inhibitor solution (Nacalai Tesque) and then disrupted with an ultra-sonic disruptor (model VCX-500, Sonics and Materials. Inc) on ice for 30 min. The supernatant fractions were collected by centrifugation at 10,000 x g at 4°C for 20 min. The sample was incubated at 75°C for 30 min and denatured proteins were removed by centrifugation at 10,000 x g at 4°C for 20 min. The supernatant fraction was loaded on to a HiTrap Q HP (5 ml, Cytiva) column. The unbound proteins were washed out with 25 mL buffer A. The bound proteins were eluted by a linear gradient of KCl from 200 mM to 1000 mM in buffer A. The fractions containing TK1257 gene product were combined and then loaded onto a HiTrap Heparin HP column (5 mL, Cytiva). The unbound proteins were washed out with 25 mL buffer A. TK1257 gene product was eluted with a linear gradient of KCl from 200 mM to 1000 mM in 50 mL buffer A. The purity of the fractions was assessed by 10% SDS-polyacrylamide gel electrophoresis (PAGE). The TK1257 gene product fractions were combined and then the buffer was changed to storage buffer [50 mM Tris-HCl (pH 7.6), 5 mM MgCl_2_, 500 mM KCl, 6 mM 2-mercaptoethanol, and 50% (v/v) glycerol] using a Vivaspin 15R centrifugal filter unit (code VS15RH01, 10,000 MWCO, Sartorius). The purified TK1257 gene product was concentrated to 3.5 mg/mL in a Vivaspin 15R centrifugal filter unit and stored at - 30°C.

### Preparation of tRNA transcripts

Transfer RNA transcripts were prepared by *in vitro* transcription using T7 RNA polymerase as reported (87). The sequences of oligonucleotides used for the construction of template DNA are given in Supplementary Table 2. The transcripts were purified on 10% polyacrylamide gels containing 7 M urea electrophoresis [10% PAGE (7 M urea)] (88).

### Measurements of tRNA methyltransferase activity of TK1257 gene product

The time course experiment was performed as follows. 3 µM TK1257 gene product, 35 µM *T. kodakarensis* tRNA^Trp^ transcript and 100 µM ^3^H-labeled SAM were incubated at 75°C in 200 μL of buffer B [50 mM Tris-HCl (pH 7.6), 5 mM MgCl_2_, 300 mM KCl and 6 mM 2-mercaptoethanol]. At 0, 1, 3, 5, 10, 20, 30 and 40 min time periods, 20 μL of the reaction mixture was spotted onto a Whatman 3MM filter. The filters were washed with 5% (w/v) trichloroacetic acid eight times at 4 °C for 5 min and then dried. Methyl group incorporation was measured by a liquid scintillation counter. This experiment was independently replicated three times (n = 3). The data was fitted to the single-exponential equation (eq. 1)

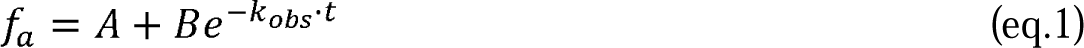

where *f_a_* is the methyl group incorporation activity [dpm] at a specific time, A is the maximum methyl group incorporation activity at completion, -B is the amplitude of the observable phase, *k_obs_* is the observed first-order rate constant for the methyl group transfer reaction, and t is time.

### Modified nucleotide analyses by two-dimensional thin layer chromatography

The reaction was performed at 75°C in 200 μL of buffer B containing 3 µM TK1257 gene product, 35 µM *T. kodakarensis* tRNA^Trp^ transcript and 100 µM ^14^C-labeled SAM. The RNA was treated with phenol-chloroform and then precipitated by ethanol. The tRNA was dissolved in 5 µl of 50 mM ammonium acetate (pH 5.3) and digested with 2.5 units of nuclease P1. 1 µl of standard nucleotide mixture (pA, pG, pC and pU, 0.05 A260 units each) was added into the sample and then spotted onto a cellulose thin-layer plate (code 1.05565.0001, Merck). The solvent systems were as follows: first dimension; isobutyric acid/concentrated NH_4_OH/water = 66/1/33 (v/v/v), second dimension; isopropyl alcohol/HCl/water = 70:15:15 (v/v/v) (89). The standard nucleotides were marked using a pencil with irradiation of ultra-violet light at 254 nm. The autoradiogram of the thin-layer plate was obtained using Typhoon FLA 7000 Laser Scanner (GE Healthcare).

Construction of T. kodakarensis ΔtrmTS strain

To construct the gene disruption plasmid (pUTS) for the *ΔtrmTS* strain, the *trmTS* (TK1257) gene along with their 5′- and 3′-flanking regions (∼1.0 kbp) were amplified from *T. kodakarensis* KUW1 genomic DNA by PCR using the primer sets TS-F/TS-R. Each fragment was inserted into the linearized plasmid pUC19 at the 5′- and 3′ blunt ends of partial BamHI sites by the in-fusion reaction. Inverse PCR was conducted with the primer sets TS-delF/TS-delR to remove the target gene (*trmTS*) from the resulting plasmid, and then the fragment was in-fused with the fragment including the PpadD-*pdaD* (Tk0149) marker gene, which was amplified with the primer set TS-AGF/TS-AGR from pTK02 plasmid (90). The pUTS plasmid was used to transform the *T. kodakarensis* KUWA strain, exhibiting agmatine auxotroph. Cells grown in MA-YT-S^0^ medium containing 1.25 mg/mL agmatine at 85 °C for 10 h were harvested and suspended in 200 μl of 0.8 × MA, and kept on ice for 30 min. 3 μg of the plasmid was gently added into the suspended cells and then kept on ice for 1h. The cells were cultivated in MA-YT-S^0^ without agmatine at 85 °C for 10 h. 100 μL of the cultures were spread onto MA-YT solid medium without agmatine and then incubated at 85 °C for 24 h. Only cells with a phenotype exhibiting agmatine prototrophy by homologous recombination (Supplementary Figure S1) can grow in the absence of agmatine. Single colonies were picked up and then cultured in MA-YT-S^0^ medium without agmatine at 85°C for 10 h. The candidate *ΔtrmTS* cells were harvested and suspended in distilled water. Genomic DNA was extracted from the cells by phenol-chloroform treatment. DNA sequencing of the recombination region was performed. The primers used for the disruption of *trmTS* gene are listed in Supplementary Table 3.

### Western blotting

Customized rabbit anti-TrmTS serum was prepared by Kitayama Labes Co., LTD, Japan. The polyclonal antibody fraction was prepared using a MAbTrap Kit (GE Healthcare). Cell extracts from *T. kodakarensis* wild-type and *ΔtrmTS* strains were prepared as follows. Wet cells (0.1 g) were suspended in 100 μL of 2x SDS-PAGE loading buffer [100 mM Tris-HCl (pH 6.8), 200 mM dithiothreitol, 2.5% SDS, 0.2% bromophenol blue, and 20% (v/v) glycerol], disrupted by sonication and boiled for 5 min. 0.5 μL of the samples were loaded onto a 12.5% SDS-polyacrylamide gel. Half of the gel was stained with Coomassie Brilliant Blue for visualization and half of the gel was electroblotted onto a polyvinylidene difluoride membrane (code 10600029, 0.45 μm, GE Healthcare). TrmTS was detected using horse radish peroxidase (HRP)-conjugated Affinipure Goat Anti-Rabbit IgG (H+L) (code AB-2313567, Proteintech) as a secondary antibody and the chemiluminescence derived from HRP was detected with a Typhoon FLA 7000 Laser Scanner (GE Healthcare) in accordance with the manufacturer’s instruction.

### Purification of tRNA^Trp^ from *T. kodakarensis* wild-type and *ΔtrmTS* strains using a solid-phase DNA probe method

Total RNAs were prepared from *T. kodakarensis* wild-type and *ΔtrmTS* strains as described previously (91). The tRNA fraction was further purified by 10% PAGE (7 M urea). Transfer RNA^Trp^ molecules were purified from the tRNA mixtures using a solid-phase DNA probe method (91, 92). The sequence of DNA probe was complimentary to U20-A36 in *T. kodakarensi*s tRNA^Trp^: 5′-TGG AGC CCG CGA TGA TGG A-biotin-3′.

### Mass spectrometry for RNA modification analysis

To generate RNase T1 fragments of RNA, 40 pmol of tRNA was digested in a 20 μL mixture containing 100 units of RNase T1 (Thermo Fisher Scientific) and 5 mM ammonium acetate (NH_4_OAc, pH 5.3) at 37°C for 60 min. To generate RNase A fragments of RNA, 40 pmol of tRNA was digested with 1 μg of RNase A (Thermo Fisher Scientific) and 1 unit of calf intestinal alkaline phosphatase (NEB), if required, in total 20 μL of 5 mM ammonium bicarbonate (AMBIC) at 37°C for 60 min. The nucleases were removed by phenol-chloroform extraction as follows: Twenty µl of water-saturated phenol was added to the reaction mixture and vigorously vortexed for 10 s, and then 20 µl of chloroform were added and vortexed as well. After centrifugation at 15,000 ×g for 5 min at room temperature, the upper aqueous layer was transferred to a new tube and dried in vacuum. The dried samples were dissolved with 10 μL of 70–80% acetonitrile (ACN) and analyzed on a liquid chromatography-mass spectrometry (LC-MS) system (Vanquish Flex UHPLC system and Orbitrap Exploris 240 (Thermo Fisher Scientific)). The digests were separated on an iHILIC-Fusion column (PEEK, 100×2.1 mm, 3.5 μm, HILCON) with a guard column (PEEK, 20×2.1 mm, 5 μm, HILCON) in 5 mM NH_4_OAc (pH 5.3) with a linear gradient of acetonitrile (80 to 50% ACN over 20 min) at 150 μl/min. Negative ion scanning ranged from 600 to 2000 *m/z*. Nucleoside measurement was conducted according to the literature (93) with slight modifications. Twenty pmol of tRNA was digested by the following 3-step reaction at 37°C for 1 h in each step: (1) 0.03 units of nuclease P1 (Fujifilm Wako Pure Chemical) in 10 mM NH_4_OAc (pH 5.3), (2) 0.04 units of phosphodiesterase I (Worthington Biochemical) in 50 mM AMBIC and (3) 0.03 units of alkaline phosphatase (*E. coli* C75, Nippon Gene) in 50 mM AMBIC. Enzymatic digests were separated on a SunShell C18 column (150×2.1 mm, 2.6 μm, ChromaNik Technologies) with a guard cartridge column RP (ChromaNik Technologies). The solvent system comprised 5 mM NH_4_OAc (pH 5.3) and ACN with a multi-step gradient (1–11% ACN from 0 to 10 min, 11–24% ACN from 10 to 20 min, 24–90% ACN from 20 to 25 min, and 90% ACN from 25 to 35 min, followed by 1% CAN equilibration for 10 min) at 100 μl/min. Positive ion scanning ranged from 105 to 700 *m/z*. Mass spec data were analyzed using the Xcalibur Qual Browser (Thermo Fisher Scientific).

### Measurements of kinetic parameters of TrmTS for SAM and tRNA

The kinetic parameters for SAM and tRNA were determined by a custom-made Python program (94), in which the data were fitted to the Michaelis-Menten equation. The following mixtures were used for the measurement of kinetic parameters of the wild-type TrmTS (or TrmTS mutant protein) for SAM: 3 µM TrmTS, 35 µM tRNA, various concentrations (0.25, 0.5, 1.0, 2.0, 4.0, 8.0, 16.0, 32.0, 64.0 µM) of SAM (a mixture of non-radioisotope and ^3^H-labeled SAM; molecular ratio 1000:1) in 25 µL of buffer B. In the case of measurement of kinetic parameters for tRNA transcripts, 3 µM TrmTS (or TrmTS mutant protein), various concentrations (0.25, 0.5, 1.0, 2.0, 4.0, 8.0, 16.0, 32.0 µM) of tRNA transcript, and 100 µM SAM were mixed in 25 μL of buffer B. The reaction mixtures were incubated at 75°C for 10 min and then spotted onto Whatman 3MM filters. Incorporation of ^3^H-methyl groups was monitored by conventional filter assay. In brief, the filters were washed in 50 mL 5% (w/v) trichloroacetic acid solution five times. The filters were dipped into 50 mL 99.5% ethanol to remove water and then dried. The ^3^H-methyl group incorporation was measured by a liquid scintillation counter. The filter assay was independently replicated three times (n = 3).

### Alignment analysis

TrmTS homologs were searched using BLAST with default algorithm parameters except for Max target sequences (increased to 500). To depict the distribution of conserved amino acids, sequence alignments were generated by ClustalW (95) and ESPript 3 (96) with sequences from eight selected organisms including *Thermococcus kodakarensis* with E-value less than 1e-20.

### Site-directed mutagenesis and purification of mutant proteins

Site-directed mutagenesis was performed using reverse polymerase chain reaction with the primers listed in Supplementary Table 4. These mutations were verified by DNA sequencing. All TrmTS mutant proteins were expressed and purified using the same methods for the wild-type TrmTS protein.

### Measurements of CD-spectra

TrmTS wild-type and mutant proteins were dialyzed against CD buffer [50 mM Tris-HCl (pH 7.6), 5 mM MgCl_2_, 500 mM KCl]. After dialysis, the protein concentrations were adjusted to 2 µM. CD spectra were measured on a JASCO J-820 spectropolarimeter equipped with a JASCO PTC-423L thermo-controller at 25, 50 and 75°C. Cuvettes with a 1 mm path length were used. The spectra were recorded from 300 nm to 200 nm. The scan speed was 50 nm/min. The spectra shown in this report are the average of five scans.

### Inhibition experiments with mutant tRNA^Trp^ transcripts

3 µM TrmTS, 35 µM the wild-type tRNA^Trp^ transcript, various concentrations (0, 10, 20, 30, 40, 50, 60, 70, 80 µM) of mutant tRNA^Trp^ transcript, 100 µM ^3^H-labeled SAM in 25 μL of buffer B were incubated at 75°C for 25 min and then the filter assay was performed as described in “Measurements of tRNA methyltransferase activity of TK1257 gene product” section. The assay was independently replicated three times (n = 3).

### Structural models

The structural model of *T. kodakarensis* TrmTS was constructed by AlphaFold 3 (97). Electro potential map was generated by APBS Electrostatics in PyMOL using default settings (−5.000 kBT/e ∼ 5.000 kBT/e). The conserved amino acid residues map was generated by Consurf-DB using default settings (98).

### Prediction of the secondary structure of truncated tRNA mutants

The secondary structure of truncated tRNA mutant transcripts (transcript 6, transcript 7, and transcript 8) were predicted using the RNAfold Web Server (http://rna.tbi.univie.ac.at//cgi-bin/RNAWebSuite/RNAfold.cgi) with default parameters.

## RESULTS

### Search for a candidate gene using comparative genomics

Initially, we searched for a candidate gene for Cm6 formation in *T. kodakarensis* tRNA by comparative genomics (Fig. 1B and C). 2,306 genes are encoded in the *T. kodakarensis* genome (99). Fortunately, activities of tRNA modification enzymes in a cell extract of *Pyrococcus furiosus* have been reported (100): when *Haloferax volcanii* tRNA^Ile^ transcript was used as a substrate, Am6 was produced by the cell extract of *P. furiosus*. The 2′-*O*-methyltransferase responsible for the Am6 formation in *P. furiosus* has not yet been identified. If our target methyltransferase catalyzes the 2′-*O*-methylation of A6 as well as C6, an ortholog of the target methyltransferase will be encoded in the *P. furiosus* genome. Therefore, we predicted that the candidate gene is encoded in both *T. kodakarensis* and *P. furiosus* genomes. In contrast, tRNAs from *H. volcanii*, *Sulfolobus acidocaldarius*, and *Thermoplasma acidophilum* do not contain the Cm6 modification (101–105). We therefore predicted that these archaea genomes do not contain the candidate gene. Using this analysis, we were able to disregard a large fraction of the *T. kodakarensis* genome such that only 570 genes remained. Of the 570 genes, ten genes are annotated as RNA methyltransferase or tRNA-binding protein genes and two genes are annotated as function unknown genes (Supplementary Table 5). One gene (TK1257) product, which is composed of 355-amino acid, is predicted to possess N-terminal ferredoxin-like, THUMP and SPOUT RNA methyltransferase domains. As described in the Introduction, the THUMP domain recognizes the CCA terminus of tRNA. Furthermore, the majority of the SPOUT RNA methyltransferase superfamily are 2′-*O*-methyltransferases (11). Moreover, the N-terminal ferredoxin-like domain often adjusts the distance and angle between the THUMP and the Rossmann fold RNA methyltransferase catalytic domains (79). Therefore, we considered the TK1257 gene to be the best candidate.

### Recombinant TK1257 gene product possesses tRNA methyltransferase activity for Cm6 formation

The TK1257 gene product was expressed in *Escherichia coli* cells and purified as shown in Fig. 2A. We observed that TK1257 gene product aggregated during dialysis against buffer with low concentrations of KCl (below 50 mM). Initially, therefore, we investigated the optimum KCl concentration for methylation by TK1257 gene product (Supplementary Fig. 1). In this experiment, we used *E. coli* tRNA mixtures because we assumed that methylation by the TK1257 gene product may require other modifications of tRNA. It should be mentioned that *E. coli* tRNAs do not possess the Cm6 modification. As shown in Supplementary Fig. 1, the velocity of methyl-group incorporation into *E. coli* tRNA mixture was maximum at 300 mM KCl. Therefore, 300 mM KCl was added to the reaction mixtures in this study. When TK1257 gene product, *T. kodakarensis* tRNA^Trp^ transcript, and ^3^H-labeled SAM were incubated at 75°C, ^3^H-methyl groups were clearly incorporated into tRNA^Trp^ transcript (Fig. 2B). Thus, this result shows that methylation by TK1257 gene product does not require any other modifications in tRNA. To address the modification position, we prepared three mutant tRNA^Trp^ transcripts in addition to the wild-type tRNA^Trp^: the C6-G67 base pair was replaced by A6-U67, U6-A67 or G6-C67 base pairs (Fig. 2C). The mutant tRNA^Trp^ A6-U67 and U6-A67 transcripts were methylated by TK1257 gene product as well as the wild-type tRNA^Trp^ transcript (Fig. 2D). In contrast, the mutant tRNA^Trp^ G6-C67 transcript was not methylated (Fig. 2D). To confirm the modified nucleotides, these tRNA^Trp^ transcripts were treated with TK1257 gene product and ^14^C-labeled SAM, and then digested with nuclease P1. The resultant nucleotides were separated by two-dimensional thin-layer chromatography and the autoradiograms of the thin-layers were obtained (Fig. 2E). As shown in Fig. 2E, the spot corresponding to ^14^C-pCm was clearly detected from the wild-type tRNA^Trp^ sample. Furthermore, ^14^C-pAm and ^14^C-pUm were detected from the tRNA^Trp^ A6-U67 and tRNA^Trp^ U6-A67 mutant samples, respectively (Fig. 2E). In contrast, no spot was observed from the tRNA^Trp^ G6-C67 mutant sample (negative control) (Fig. 2E) consistent with the results in Fig. 2D. Thus, these results clearly show that the methylation position is the ribose at position 6 in tRNA. Furthermore, the result with tRNA^Trp^ A6-U67 mutant transcript is in line with the observation in an early study (100): when *H. volcanii* tRNA^Ile^ transcript was used as a substrate, Am6 was formed by *P. furiosus* cell extract. The modification position was further checked by LC/MS analysis of methylated tRNA^Trp^ (Supplementary Fig. 2) and tRNA^Pro^ (Supplementary Fig. 3) transcripts. When the ribose at position 6 in tRNA^Trp^ transcript was methylated by TK1257 gene product, RNase A did not cleave the phosphodiester bond between C6 and G7. As a result, an RNA fragment, 5′-HO-GGGGGCmGUp-3′ (m/z 876.79) appeared (Supplementary Fig. 2). Similarly, when methylated tRNA^Pro^ was digested with RNase T1, 5′-HO-CCmGp-3′ (m/z 986.13) appeared (Supplementary Fig. 3). Taking these experimental results together, we conclude that the TK1257 gene product possesses a tRNA methyltransferase activity for Cm6 formation in tRNA^Trp^ and tRNA^Pro^ transcripts. Based on these results, hereafter we describe TK1257 gene product as TrmTS (<u>T</u>ransfer <u>R</u>NA <u>m</u>ethylation gene product with <u>T</u>HUMP and <u>S</u>POUT domains). To investigate the enzymatic properties of TrmTS, kinetic parameters of TrmTS and tRNA^Trp^ transcript were determined at 75°C. Because the Km value of TrmTS for the wild-type tRNA^Trp^ transcript was determined to be 7.1 μM (Fig. 2F), we used 100 μM RNAs as substrates in latter experiments. Furthermore, kinetic analyses of mutant tRNA^Trp^ A6-U67 and U6-A67 transcripts reveal that these mutant tRNA^Trp^ transcripts are efficiently methylated by TrmTS. It should be noted that all tRNA molecular species encoded in the *T. kodakarensis* genome possess C6 or G6 (98). Therefore, the Am6 and Um6 formations by TrmTS do not occur in native tRNAs. Moreover, the Km value of TrmTS for SAM was determined to be 20.3 μM (Fig. 2G). This Km value is slightly large for a Km value of tRNA methyltransferases for SAM: for example, the Km value of *T. thermophilus* TrmH for SAM is 10.0 μM (26). Based on this experimental result, we used 200 μM SAM as the methyl group donor in latter experiments.

**Figure 2.**
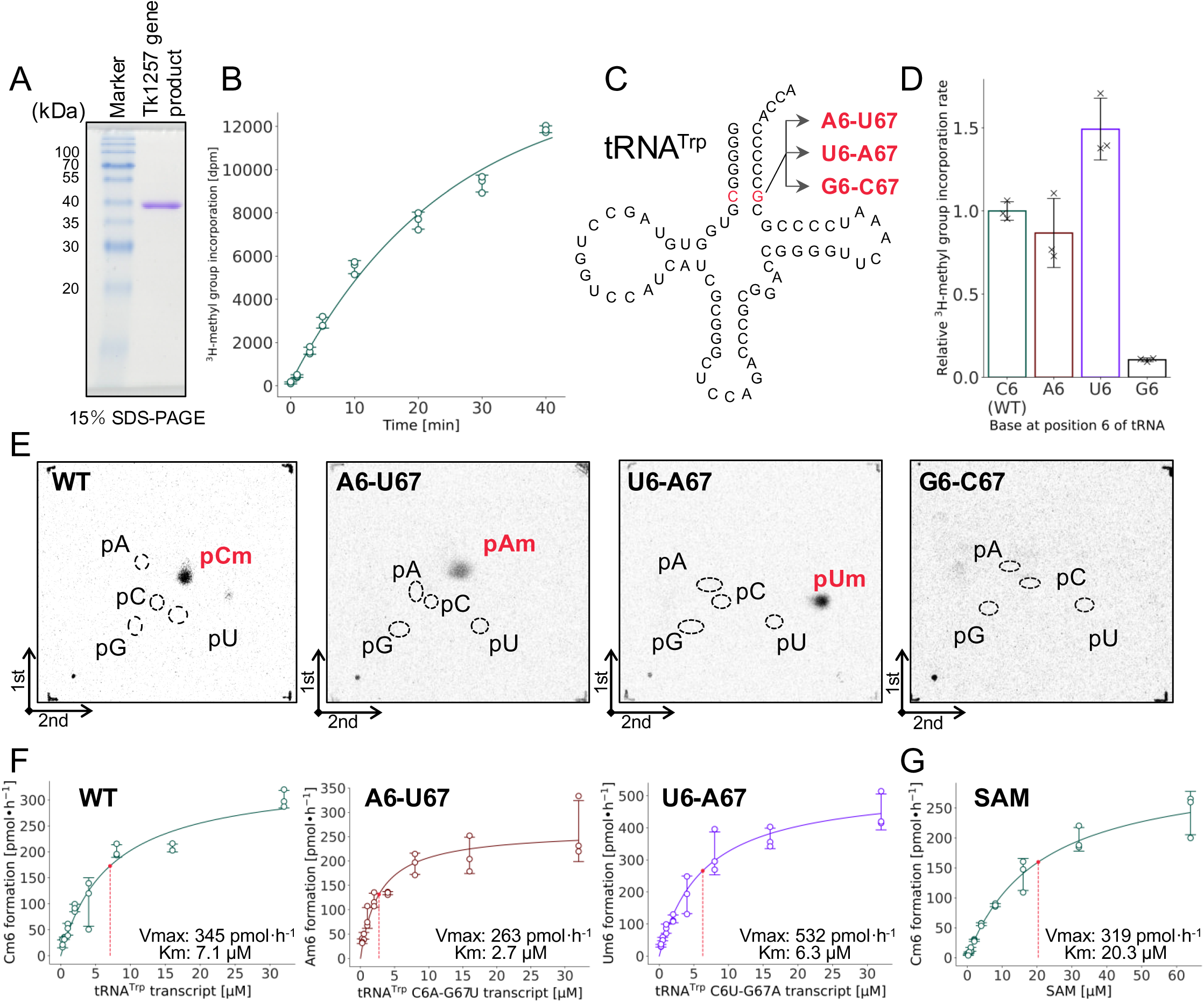
TK1257 gene product possesses a tRNA methyltransferase activity for 2′-*O*-methylation at position 6. A, Purified TK1257 gene product (3 μg) was analyzed by 15% SDS-PAGE. The gel was stained with Coomassie Brilliant Blue. B, The methyl-transfer activity of TK1257 gene product was tested at 75°C using tRNA^Trp^ transcript and ^3^H-labeled SAM. This experiment was independently replicated three times (n = 3). The error bars show the standard deviations. C, Three mutant tRNA^Trp^ transcripts, in which the C6-G67 base pair was replaced by A6-U67, U6-A67 or G6-C67 base pair, were prepared. The mutation site is highlighted in red. D, The methylation speeds for the wild-type and mutant tRNA^Trp^ transcripts were compared. The methylation for the wild-type tRNA^Trp^ is expressed as 1.00. This experiment was independently replicated three times (n = 3). The error bars show the standard deviations. E, ^14^C-methylated nucleotides were analyzed by two-dimensional thin-layer chromatography. When the wild-type tRNA^Trp^ transcript (WT) was used, ^14^C-pCm was detected. In contrast, when the mutant tRNA^Trp^ A6-U67 and U6-A67 transcripts were used, ^14^C-pAm and ^14^C-pUm were detected, respectively. Furthermore, when the mutant tRNA^Trp^ G6-C67 transcript was used, ^14^C-labeled nucleotide was not detected, consistent with the result in panel D. F, Kinetic parameters of TK1257 gene product for the wild-type and mutant tRNA^Trp^ A6-U67 and U6-A67 transcripts were determined. These transcripts had comparable methyl group acceptance activities. G, Kinetic parameters of TK1257 gene product for SAM were determined.

### The *ΔtrmTS* strain showed a slight growth retardation at 93°C

To confirm that the *trmTS* gene is expressed in living cells, we constructed a *trmTS* gene deletion (*ΔtrmTS*) strain (Supplementary Fig. 4). To detect TrmTS, we prepared anti-TrmTS polyclonal antibody fraction from an immunized rabbit. Proteins in the cell-extracts of the wild-type and *ΔtrmTS* strains were separated by 12.5% SDS-polyacrylamide gel electrophoresis (PAGE) (Fig. 3A). 20 ng of purified TrmTS was used as a positive control (right lane in Fig. 3A): the band of TrmTS is not visible by Coomassie Brilliant Blue staining because only 20 ng of protein was used. The proteins were electro-blotted onto a polyvinylidene difluoride membrane filter and then western blotting analysis was performed using the anti-TrmTS polyclonal antibody fraction. The band corresponding to the TrmTS was clearly observed in the cell-extract of the wild-type strain. In contrast, this band was not observed in the cell-extract of the *ΔtrmTS* strain. This result clearly shows that TrmTS is expressed in the wild-type strain and this protein is not expressed in the *ΔtrmTS* strain. Transfer RNA^Trp^ molecules were purified from the wild-type and *ΔtrmTS* strains by the solid-phase DNA probe method (91, 92) (Fig. 3B). The methylated cytidines in these tRNA^Trp^ molecules were analyzed by LC/MS (Fig. 3C). As shown in Fig. 1A, *T. kodakarensis* tRNA^Trp^ possesses two m^5^C and four Cm modifications. In the tRNA^Trp^ from the wild-type strain, the amount of Cm is larger than that of m^5^C (Fig. 3C left). In contrast, in the tRNA^Trp^ from the *ΔtrmTS* strain the amount of Cm is smaller than that of m^5^C due to disappearance of Cm6 (Fig. 3C right). It should be mentioned that the modification level of each Cm modification is not 100%. Therefore, this data shows decrease only in the total Cm modification level. Taking these experimental results together, we conclude that TrmTS is expressed in living *T. kodakarensis* cells and methylates tRNA^Trp^ *in vivo*. To address the phenotype of the *ΔtrmTS* strain, we compared the growth curves of the wild-type and *ΔtrmTS* strains at 85 and 93°C. As shown in Fig. 3D, the *ΔtrmTS* strain showed a slight growth retardation at 93°C: doubling times of the wild-type and *ΔtrmTS* strains were 2.19 h and 2.58 h, respectively. When TrmTS was expressed in the *ΔtrmTS* strain, the growth phenotype was recovered (Fig. 3D right panel, blue line). Therefore, the presence of TrmTS may help the survival of *T. kodakarensis* at high temperatures.

**Figure 3.**
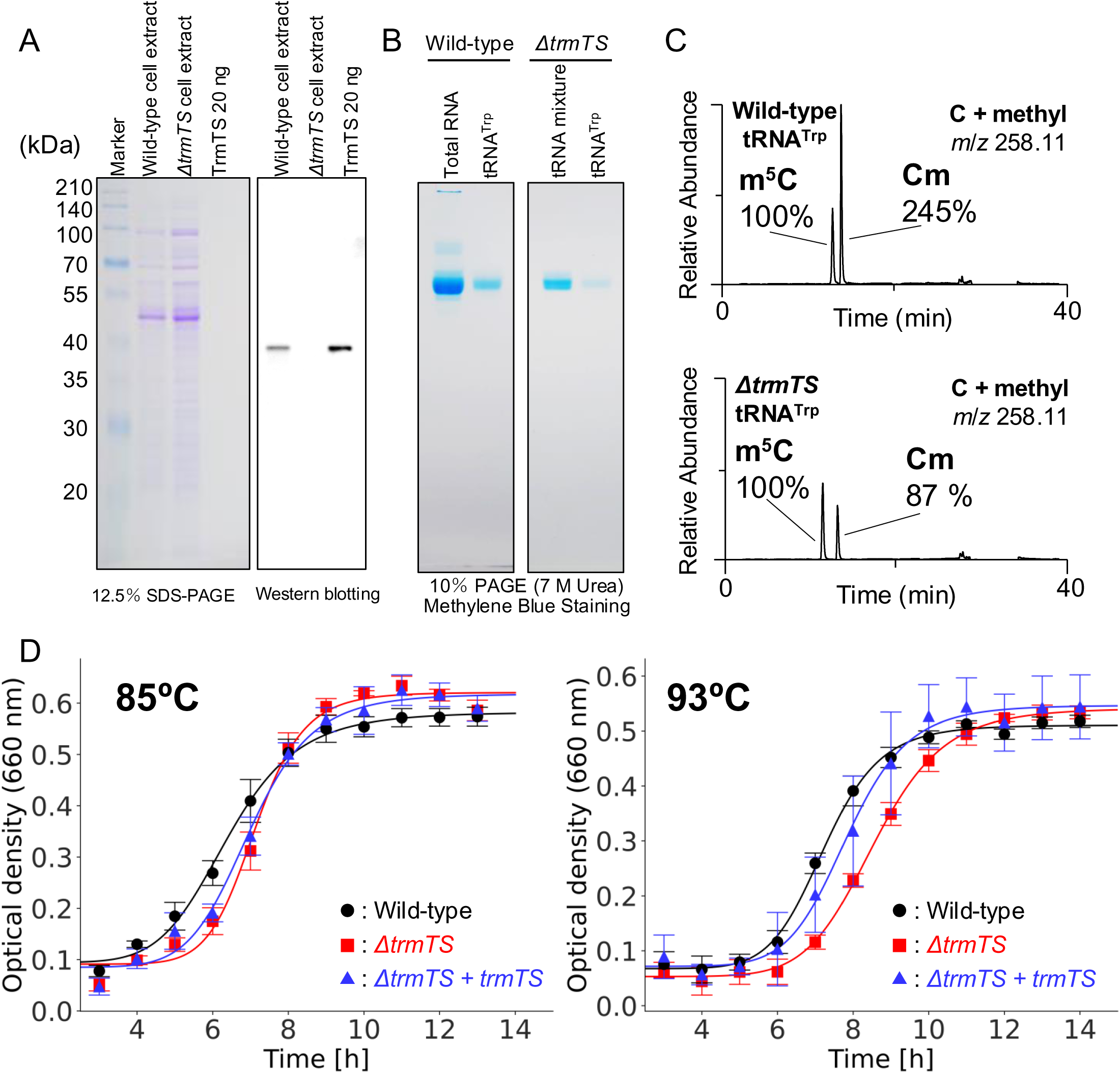
TrmTS is expressed in *T. kodakarensis* wild-type strain and the *trmTS* gene deletion (Δ*trmTS*) strain showed a slight growth retardation at 93°C. A, The cell extracts from wild-type and *ΔtrmTS* strains were analyzed by western-blotting. Purified TrmTS (right lane) was used as a positive control. The gel was stained with Coomassie Brilliant Blue. Because only 20 ng of TrmTS was used, the band of TrmTS was not visible by Coomassie Brilliant Blue staining. **B**, Native tRNA^Trp^ were purified from the wild-type and *ΔtrmTS* strains. The gel was stained with methylene blue. **C**, Contents of methylated cytidines (m^5^C and Cm) in purified tRNA^Trp^ from the wild-type (upper) and *ΔtrmTS* (lower) strains were analyzed by LC/MS. The peak area of the m^5^C in tRNA^Trp^ from the wild-type strain is expressed as 100%. **D**, Growth curves of the wild-type (black) and *ΔtrmTS* (red) strains are compared at 85°C and 93°C. At 93°C, a slight growth retardation of *ΔtrmTS* strain was observed. When TrmTS was expressed in the *ΔtrmTS* strain (blue), the growth speed recovered. This experiment was independently replicated three times (n = 3). The error bars show the standard deviations.

### Identification of three conserved amino acid sequence motifs and amino acid residues essential for enzymatic activity

Using the amino acid sequence of *T. kodakarensis* TrmTS as a query, we searched for TrmTS-like proteins in other organisms. As shown in Fig. 4A, TrmTS-like proteins are encoded in several thermophilic archaea and eubacteria. The amino acid sequence alignment predicted that TrmTS contains N-terminal ferredoxin-like, THUMP and SPOUT domains. Between the THUMP and SPOUT domains, a linker region exists. Also, we identified several conserved amino acid residues in these TrmTS-like proteins (Fig. 4A). TrmH is known to possess three conserved amino acid sequence motifs, namely motifs 1, 2 and 3 (9). The catalytic center is an arginine residue in motif 1 (Arg41 in TrmH) (25–27). The conserved amino acid residues in motif 2 and motif 3 contribute to the formations of the trefoil-knot structure and SAM-binding pocket (25, 26). To investigate the catalytically and structurally important residues in TrmTS, we first examined if TrmTS also possesses these motifs found in TrmH. A multiple sequence alignment of the SPOUT domain sequences from TrmTS, TrmL, TrmJ, Trm56, and TrmH revealed that motifs 1, 2, and 3, as characterized in TrmH, are not clearly conserved in TrmTS (Supplementary Fig. 5A).

**Figure 4.**
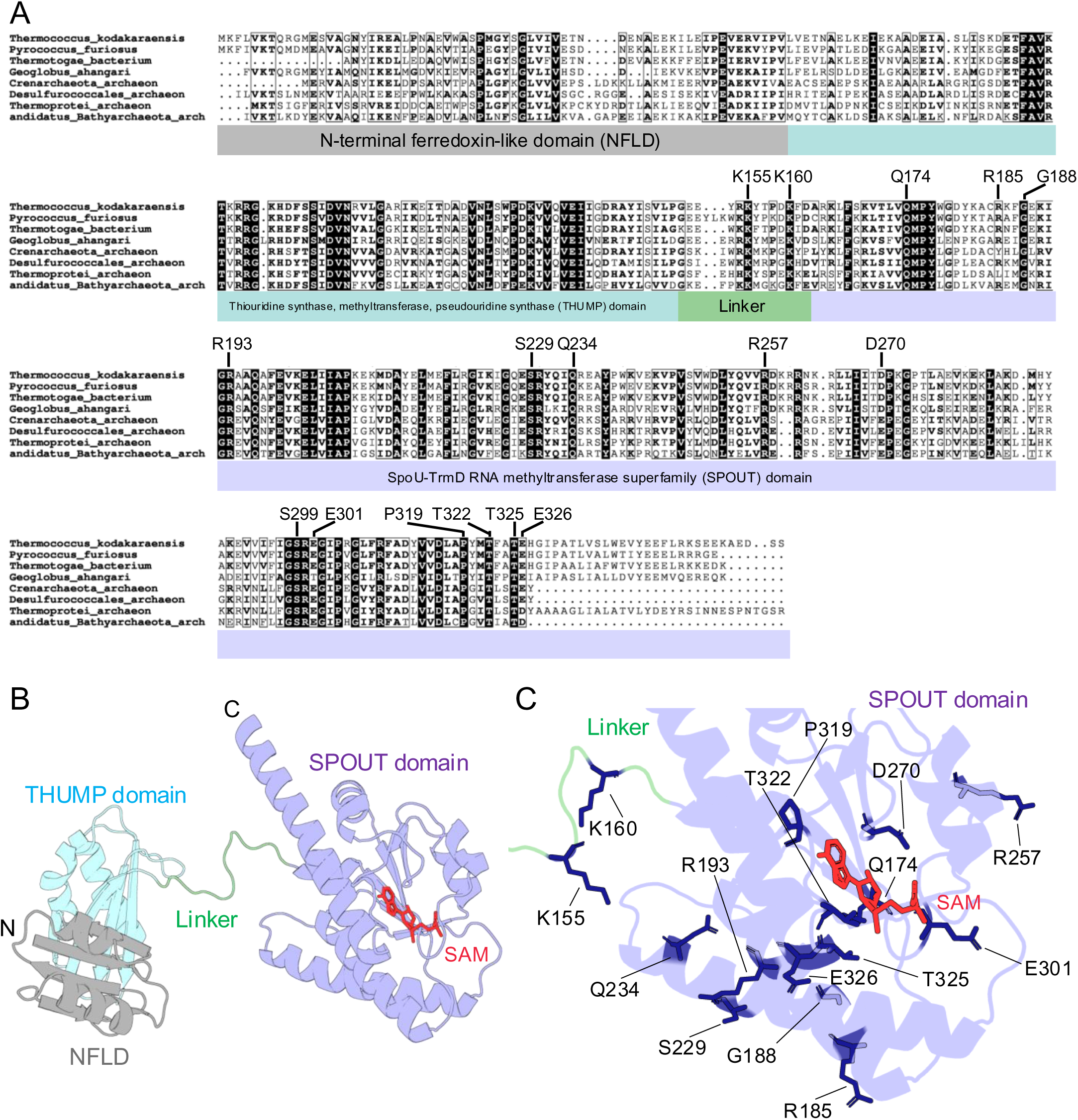
Amnio acid sequence alignment of TrmTS-like proteins and SPOUT tRNA 2′-*O*-methyltranserases and location of conserved amino acid residues. A, The amino acid sequence alignment of TrmTS-like proteins is depicted. N-terminal ferredoxin-like (NFLD), THUMP and SPOUT domains, and linker region are highlighted in gray, pale blue and purple, and green, respectively. Fifteen conserved amino acid residues were substituted by alanine and/or other amino acid residues. B, Subunit structural model of TrmTS was constructed by AlphaFold 3. A SAM molecule was modeled on the structure where the crystal structure of *T. thermophilus* TrmH in complexed with SAM (PDB: 1V2X) was aligned with the TrmTS structure. The TrmH structure was then removed. SAM is colored red. The domains and linker region are colored in the same as in panel A. C, The mutation sites are mapped onto the SPOUT catalytic domain and linker region.

Next, we constructed a structural model of TrmTS using AlphaFold 3 (Fig. 4B). The parameters of the structural model are given in Supplementary Table 6. The predicted template modeling (pTM) value of monomer subunit is 0.78: in general, the pTM value more than 0.5 means that the model has been constructed with high precision. Subsequently, the conserved amino acid residues in TrmTS were mapped onto the structural model of the SPOUT domain (Fig. 4C and Supplementary Fig. 5B). For this purpose, we constructed a dimer structure model of TrmTS by AlphaFold 3, as all previously characterized SPOUT 2′-*O*-methyltransferases are known to form a dimer structure (Supplementary Fig. 5C, Supplementary Fig. 6, and ref. 11). The ipTM value of the dimer model is 0.78, indicating a confident prediction. The location of dimer interface of the model coincided with that observed in known SPOUT 2′-*O*-methyltransferases (Supplementary Fig. 6A): the hydrophobic areas (area size, 2635.6 _^2^; orange area in the SPOUT catalytic domain of TrmTS in Supplementary Fig. 6A) of monomer subunits assemble into the dimer structure. Furthermore, topological diagrams of TrmTS and other SPOUT enzymes showed similar structural arrangements (Supplementary Fig. 6B). These findings strongly suggest that TrmTS forms a dimer structure like other SPOUT 2′-*O*-methyltransferases. To determine whether TrmTS forms a dimer structure, we analyzed the purified full-length TrmTS by Superdex-200 gel-filtration column chromatography (Supplementary Fig. 7). The protein was eluted in the fractions I-IV (Supplementary Fig. 7A and 7B). The peak observed in the fraction IV corresponds to the molecular size of TrmTS dimer: the elution points of molecular weight markers are shown by arrows in Supplementary Fig. 7A. In addition to the dimer peak, two additional peaks were observed: one peak in fraction I, which eluted at the void volume, and another peak in fractions II and III, corresponding to the molecular size of TrmTS tetramer. Thus, the full-length TrmTS formed three peaks. As shown in Supplementary Fig. 6A, the THUMP domain of TrmTS possesses hydrophobic areas. Therefore, in the absence of substrate tRNA, purified TrmTS may form a tetramer and multimer structures in addition to a dimer structure. Furthermore, because the predicted dimer structure of TrmTS is not globular (Supplementary Fig. 6A), the molecular weight of full-length TrmTS dimer is not correctly measured by gel-filtration column chromatography. To directly assess dimerization mediated by the SPOUT catalytic domain, we prepared the SPOUT catalytic domain of TrmTS lacking the N-terminal ferredoxin-like and THUMP domains (inset in Supplementary Fig. 7C). When the SPOUT catalytic domain of TrmTS was analyzed by Superdex-200 gel-filtration column chromatography, the protein was eluted at the location of dimer (molecular weight, 48 kDa) as a single peak (peak ii in Supplementary Fig. 7C). Because the peaks i and iii do not contain any protein (Supplementary Fig. 7D), this experimental result clearly shows that the SPOUT catalytic domain of TrmTS forms a dimer.

Based on these findings, we performed single-point mutational analyses. We selected several amino acid residues for mutation, based on structural comparison between TrmTS and TrmH (Supplementary Fig. 5B and Supplementary Fig. 5C). The mutant TrmTS proteins were purified as shown in Fig. 5A. The relative initial velocities of mutant proteins were assessed (Fig.5B). After these pilot experiments, we measured the CD spectra of both wild-type and mutant proteins (Fig. 6A). Furthermore, we determined the kinetic parameters of the mutant proteins (Fig. 6B and C). These experiments elucidated that mutations in several conserved amino acid residues (e.g. Lys155, Arg185, Ser229 and Glu326, as shown in Fig. 6A) altered the peak shape of CD spectra compared to the wild-type protein, suggesting that these residues are structurally important. The amino acid sequence alignment and structural comparison between TrmTS and TrmH suggests that Arg193 is the catalytic residues. Consistent with this, the R193A mutant protein lost all enzymatic activity (Fig. 5B). Furthermore, the CD spectra indicated that the substitution of this Arginine with alanine does not affect the overall protein structure (Fig. 6C). Thus, it is likely that the Arg193 residue plays a catalytic role rather than a structural one. Next, we identified the serine residue conserved in TrmH. In the case of TrmH, Ser150 in the motif 3 in one subunit forms a hydrogen bond with the catalytic center (Arg41) in another subunit and is essential for the methyl-transfer activity (26) (Supplementary Fig. 5C). Initially, we assumed that Ser299 in TrmTS might be the serine residue that plays the same role with that Ser in TrmH. However, the Ser299A mutation did not influence the enzymatic activity (Fig. 5B). We checked the structural model again and found that Thr322 could move near to the catalytic Arg193 residue because Thr322 is located in a polypeptide loop (Supplementary Fig. 5C). Therefore, we substituted Thr322 with serine or alanine. As expected, the T322A mutant protein completely lost enzymatic activity (Fig. 5B). Furthermore, the substitution of Thr322 by serine (T322S mutant) also abolished the enzymatic activity (Fig. 5B). This result shows that the methyl group of Thr322 is essential for the enzymatic activity of TrmTS. Because this residue is conserved as serine in TrmH, the methyl group of Thr322 may possess a specific role in, for example, stabilization of trefoil-knot structure in the case of TrmTS. To address this issue, we measured the CD spectra of T322A and T322S mutant proteins (Fig. 6A). The CD spectra of these mutant proteins coincide with that of the wild-type enzyme. Thus, these results suggest that the mutation of Thr322 residue does not cause the disruption of trefoil-knot structure. One possibility is that the methyl group of Thr322 is involved in SAM-binding (formation of SAM-binding pocket). To clarify the role of Thr322, structural study of the complex between TrmTS and SAM (or tRNA) is necessary. Next, we identified the glutamic acid residue conserved in TrmH. The structural model suggested that Glu301 residue in TrmTS corresponds to the glutamic acid residue in the motif 2 in TrmH. When the Glu301 residue was substituted by alanine, considerable enzymatic activity remained (Fig. 6B). This phenomenon was also reported in the mutant protein of Trm56 (27). Although the glutamic acid residue in the motif 2 is important for folding of the catalytic domain of TrmH (26), the glutamic acid residue may be not so important for the activities of archaeal SPOUT 2′-*O*-methyltransferases including Trm56 and TrmTS. The Glu326 residue in TrmTS was expected to be the amino acid residue corresponding Asn150 residue in TrmH which is essential for the TrmH activity (26). When Glu326 was substituted by alanine, the enzymatic activity of TrmTS was completely lost (Fig. 5B and Fig. 6C). Thus, Glu326 residue is one of the conserved amino acid residues in TrmTS that plays a catalytic role. The P319 residue in TrmTS was expected to be the amino acid residue corresponding to the P143 residue in TrmH. This proline residue is semi-conserved in TrmH and is replaced by glutamine in some eukaryotic SPOUT RNA 2′-*O*-methyltransferases (for example, *Saccharomyces cerevisiae* Trm3 (105)). We expressed the TrmTS P139A mutant protein in *E. coli* cells, however the expressed protein was heavily degraded in *E. coli* cells. Therefore, we could not obtain the mutant protein. This result suggests that the Pro319 residue in TrmTS is structurally important. We continued the alanine-substitution experiments and found that six amino acid residues (Arg193, Ser229, Asp270, Thr322, Thr325 and Glu326) were essential for the methyl-transfer activity of TrmTS (Fig. 6B). Because Ser229 and Thr325 assemble around the catalytic center (Arg193), these residues are probably involved in the catalytic mechanism and/or binding of substrate(s). To clarify the roles of these residues, a crystal structure study of substrate (SAM or tRNA)-bound forms is necessary: these residues probably move according to substrate-binding.

**Figure 5.**
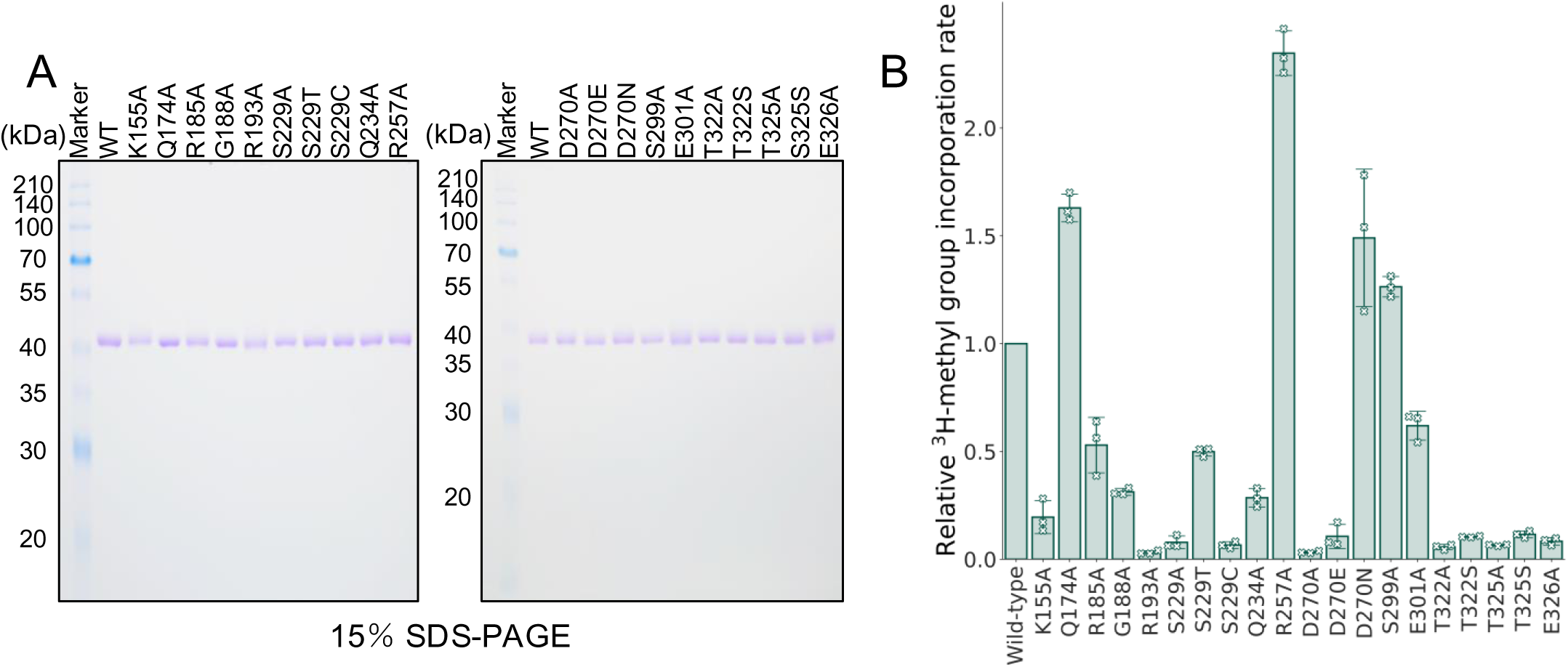
Purities of TrmTS mutant proteins and their activities. A, The wild-type and mutant TrmTS proteins (2 μg each) were analyzed by 15% SDS-PAGE. The gel was stained with Coomassie Brilliant Blue. B, Relative methylation speed of the wild-type and mutant TrmTS proteins were compared using tRNA^Trp^ transcript and ^3^H-SAM as substrates. The methylation speed of the wild-type TrmTS is expressed as 1.00. This experiment was independently replicated three times (n = 3). The error bars show the standard deviations.

**Figure 6.**
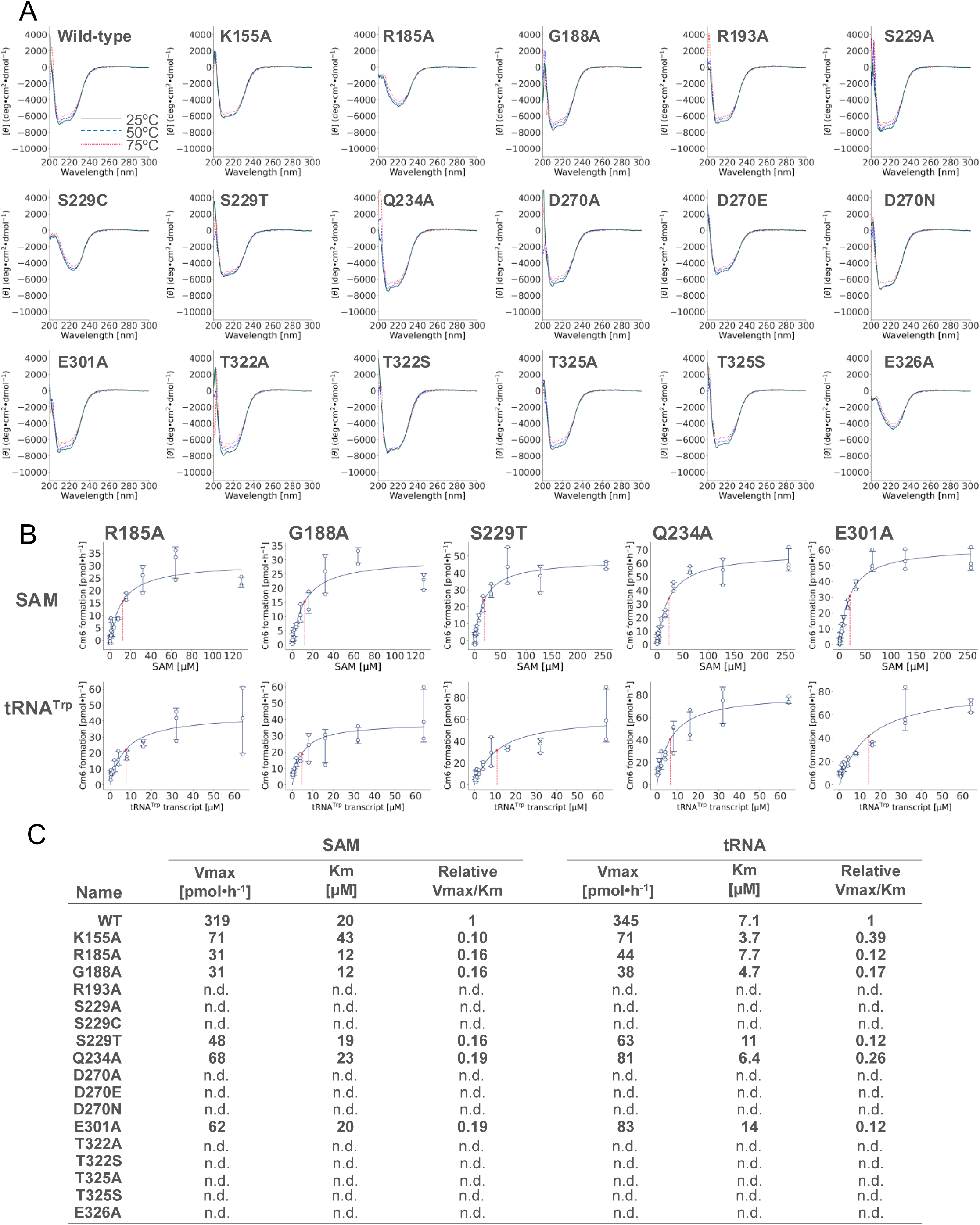
CD spectra and kinetic parameters of TrmTS mutant proteins. A, CD spectra were measured at 25, 50 and 75°C. B, Kinetic parameters of mutant TrmTS proteins for SAM and tRNA^Trp^ transcripts were measured. C, Kinetic parameters of the wild-type and mutant proteins are summarized. “n. d.” means that methyl-transfer activity was not detectable.

### The TrmTS-specific linker region contains conserved lysine residues

The multiple sequence alignment of TrmTS-like proteins showed that these proteins possess a linker region between the THUMP and SPOUT domains (Fig. 4A). This linker region is not observed in other SPOUT RNA methyltransferases. Thus, the conserved amino acid sequence in the linker region is TrmTS-specific. Two lysine residues (Lys155 and Lys160) are conserved in all TrmTS-like proteins. This linker locates near to the Asp270 residue of the catalytic domain in another subunit (Fig. 7) and the ε-amino group of Lys160 can contact the Asp270 residue. Furthermore, the replacement of Asp270 by alanine causes the complete loss of enzymatic activity. To address the role of the linker region, we substituted Lys155 and Lys160 residues with alanine. In the case of TrmTS Lys155Ala mutant protein, the methyl-transfer activity was considerably decreased (Fig. 5B). The measurement of kinetic parameters for SAM and tRNA^Trp^ transcript (Fig. 6B and C) revealed that the substitution of Lys155 with alanine critically decreases the Vmax value. Because the Lys155 residue is located far from the catalytic pocket, the Lys155 residue may be involved in movement of domains during the methylation process. The expressed TrmTS Lys160Ala mutant protein was heavily degraded in *E. coli* cells. Therefore, we could not obtain purified TrmTS Lys160Ala mutant protein. This result suggests the importance of the linker region as a structural factor.

**Figure 7.**
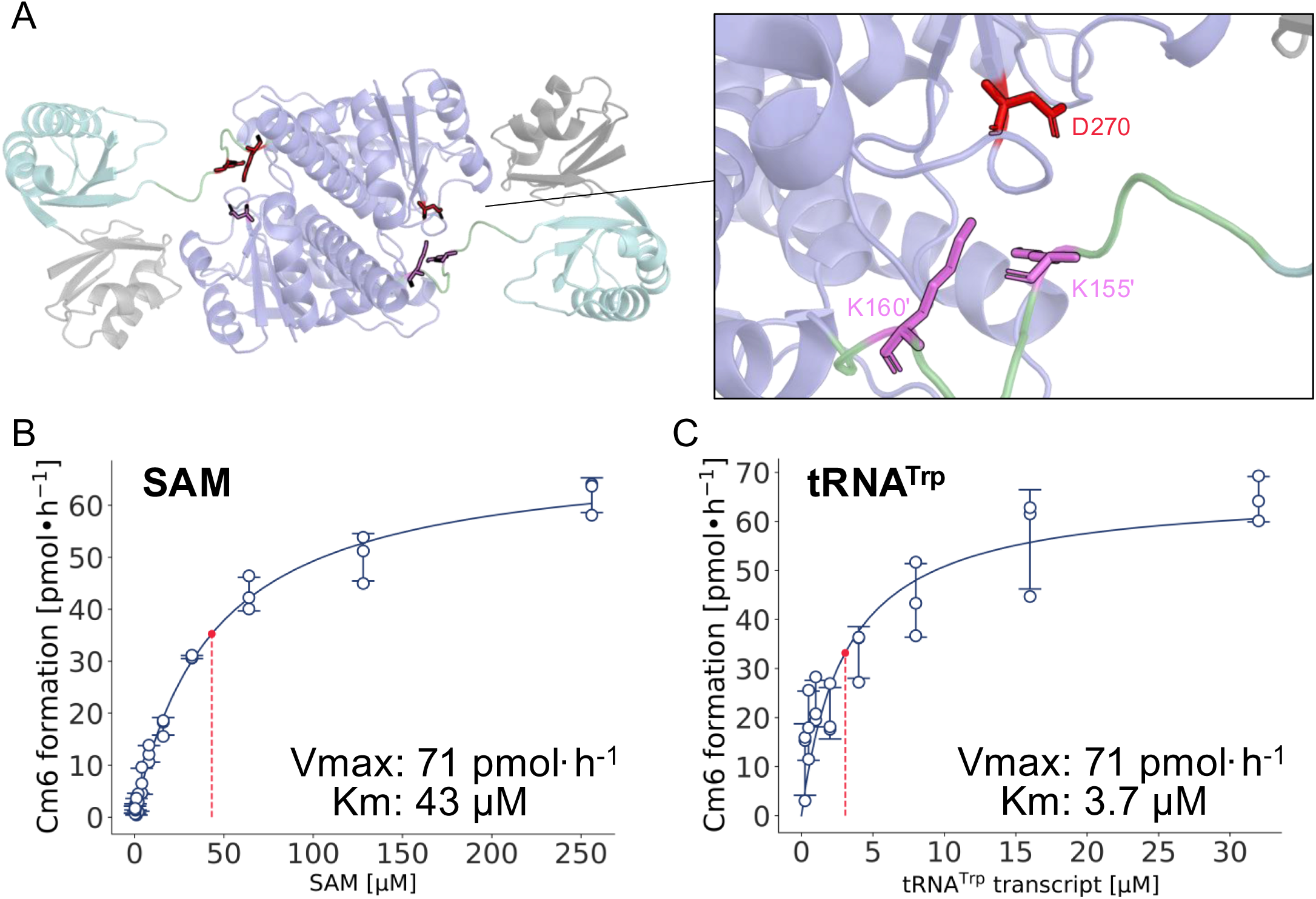
Two conserved amino acid residues (Lys155 and Lys160) in the linker region. A, The linker from one monomer contacts the SPOUT domain from the other monomer. Asp270 residue of the catalytic domain in one subunit is able to contact with Lys160 residue of the linker region in another subunit. B and C, Kinetic parameters of TrmTS Lys155Ala mutant protein for (B) SAM and (C) tRNA^Trp^ transcript were determined.

### The recognition sites of TrmTS in tRNA

To clarify the binding-mode of TrmTS to tRNA, we prepared eight truncated tRNA mutant transcripts (Fig. 8A and B). These truncated tRNA mutant transcripts were incubated with TrmTS and ^14^C-labeled SAM, and then RNAs were analyzed by 10% PAGE (7 M urea) (Fig. 8C left panels). Autoradiograms of the same gels were obtained (Fig. 8C right panels). When the D-arm (transcript 2), the anticodon-arm (transcript 3) and the T-arm (transcript 4) were individually deleted, the methyl group acceptance activities were retained (Fig. 8C). Thus, the L-shaped tRNA structure is not essential for methylation by TrmTS. In contrast, when the discriminator and CCA terminus (ACCA) were deleted (transcript 5), the methyl group acceptance activity was completely lost (Fig. 8C lane 5). Thus, the 3′-terminal region is essential for methylation by TrmTS. Furthermore, a deletion mutant of the anticodon- and T-arms (transcript 6), a deletion mutant of the D- and T-arms (transcript 7) and a deletion mutant of the D- and anticodon-arms were methylated by TrmTS (Fig.8C lanes 6-8). In contrast, a micro-helix RNA, which mimics the aminoacyl-stem, was not methylated by TrmTS at all (Fig. 8C lane 9). To explore these results deeply, we predicted the secondary structure of these truncated transcripts (Supplementary Fig. 8). The transcript 6, in which the anticodon arm and T-arm were removed, was predicted to have the acceptor stem and two stems forming a three-way junction with a loop region (Supplementary Fig. 8A). The transcript 7, in which the D-arm and T-arm ware removed, was predicted to fold into a long stem and loop structure with three short bulges (Supplementary Fig. 8B). Finally, the transcript 8 in which D-arm and anticodon arm were removed, was predicted to have an extended acceptor stem, a stretch loop, and a shortened T-arm (Supplementary Fig. 8C). Of these mutant transcripts, transcript 7 showed relatively low methyl group acceptance activity (Fig. 8C), suggesting that TrmTS recognize a minimal substrate RNA having a stretch loop connected with at least two stems for 2′-*O*-methylation, and the L-shaped tRNA structure itself is not essential.

**Figure 8.**
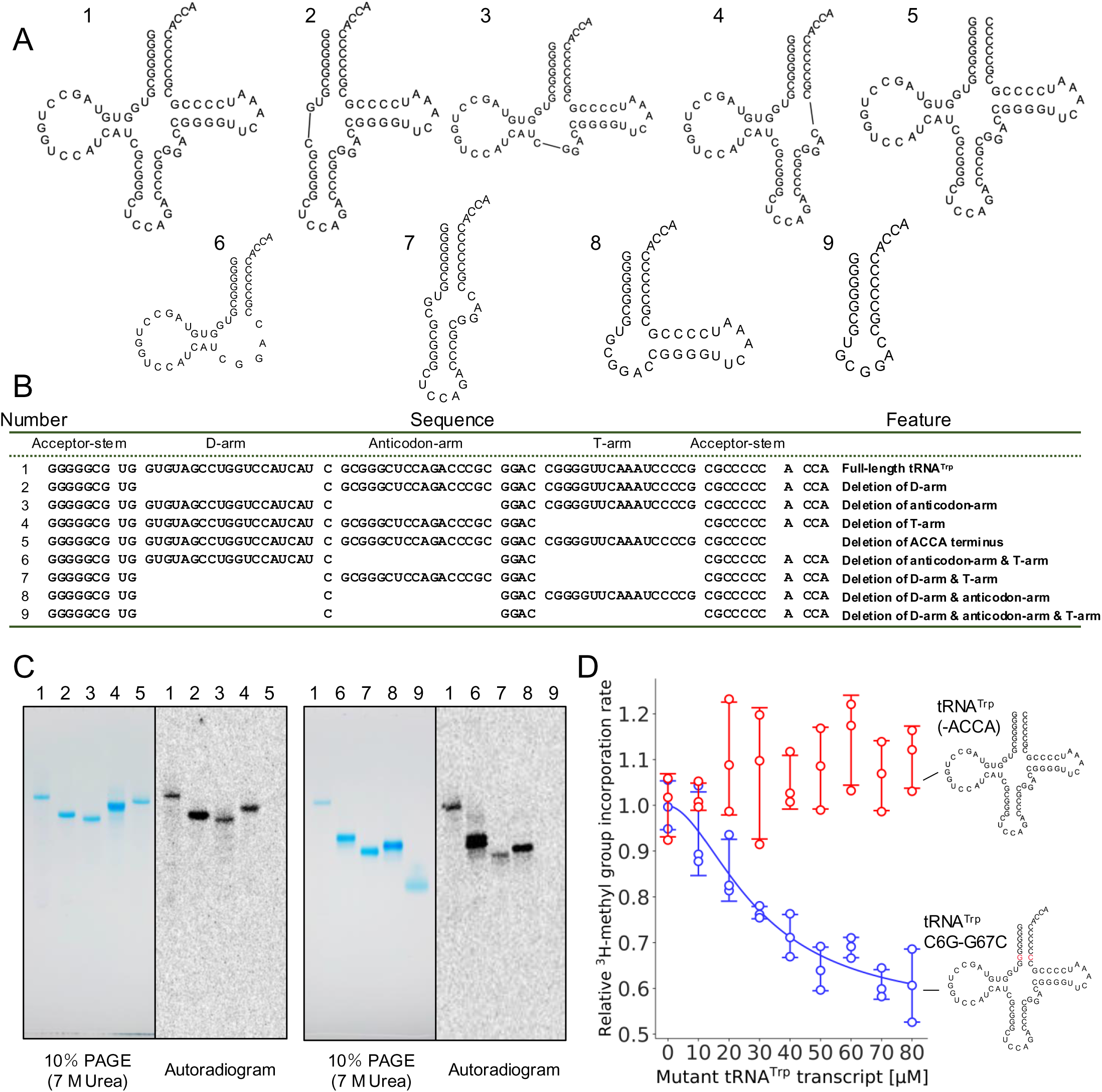
Recognition sites in tRNA. **A**, Eight mutant tRNA^Trp^ transcripts were prepared. Transcript 1 is the wild-type tRNA^Trp^ transcript. **B**, The features of mutant tRNA^Trp^ transcripts are summarized. **C**, The mutant tRNA^Trp^ transcripts were treated with TrmTS and 14C-SAM and then analyzed by 10% PAGE (7 M urea). The gel was stained with methylene blue (left). The autoradiogram of the same gel was obtained (right). The lane numbers correspond to the transcript numbers in panel A and B. **D**, Two mutant tRNA^Trp^ transcripts (-ACCA; red and G6-C67; blue) which are not methylated by TrmTS, were used for the inhibition experiments. Each mutant tRNA^Trp^ transcript (0 µM, 10 µM, 20 µM, 30 µM, 40 µM, 50 µM, 60 µM, 70 µM, and 80 µM) was titrated to the TrmTS reaction in the presence of 35 µM wild-type tRNA^Trp^. The reaction was performed at 75 °C for 10 min. The experiments were replicated three times (n=3). The error bars show the standard deviations.

### The THUMP domain of TrmTS is required for the methylation in the binding process

For TrmH, a SPOUT tRNA 2′-*O*-methyltransferase, the tRNA-binding mode consists of at least two processes, namely the initial binding and induced-fit processes (107, 108). To clarify whether the THUMP domain of TrmTS works in the binding process, we performed an inhibition experiment. Two tRNA^Trp^ mutant transcripts were prepared. One mutant tRNA^Trp^ transcript is a deletion mutant of the 3′-ACCA. Another mutant tRNA^Trp^ transcript possesses a G6-C67 base pair instead of the C6-G67 base pair. Neither mutant tRNA^Trp^ transcripts were methylated by TrmTS as shown in Fig. 2 and Fig. 8C. We tested whether these transcripts act as competitive inhibitors. When the concentration of tRNA^Trp^ G6-C67 mutant transcript was increased, the methyl-transfer to the wild-type tRNA^Trp^ transcript was clearly inhibited (blue in Fig. 8D): the IC_50_ of tRNA^Trp^ G6-C67 mutant transcript was around 30 μM. Given that the Km value of TrmTS for the wild-type tRNA^Trp^ transcript is 7.1 μM (Fig. 7C), this inhibitor possesses a considerable affinity for TrmTS. In contrast, when the concentration of tRNA^Trp^ (-ACCA) was increased, the methylation speed of the wild-type tRNA^Trp^ was not decreased (red in Fig. 8D). This result clearly shows that the CCA terminal region in tRNA is essential for the binding of TrmTS. Thus, the THUMP domain, which captures the CCA terminal region, is required in the binding process.

### Prediction of tRNA binding mode

Figure 9A shows the electro-charges of the TrmTS surface: the blue and red areas represent positive and negative charges, respectively. The amino acid residues in the positively charged areas of the THUMP and SPOUT domain are highly conserved in TrmTS-like proteins (Fig. 9B). We therefore predicted the tRNA binding mode of TrmTS based on the experimental results (Fig. 9C). When a tRNA structure was modeled onto the TrmTS structure, both tRNA and TrmTS structures were crushed. This observation suggests that the methylation by TrmTS requires an induced-fit process. It should be mentioned that clashes between the enzyme and tRNA structures is often observed in other docking models between SPOUT 2′-*O*-methyltransferases and the substrate tRNAs (38, 109, 110). Furthermore, in the case of TrmD (a member of SPOUT tRNA methyltransferases), movements of the SPOUT catalytic and C-terminal RNA binding domains according to SAM and tRNA bindings have been reported (111–113). In the binding process, the CCA terminal region is captured by the THUMP domain of TrmTS and then the three-dimensional core of tRNA contacts the positively charged areas of TrmTS. During the induced-fit process, movements of the linker region and THUMP domain may be cased. Finally, the ribose of C6 in tRNA is introduced into the catalytic pocket and accesses the catalytic Arg193 residue.

**Figure 9.**
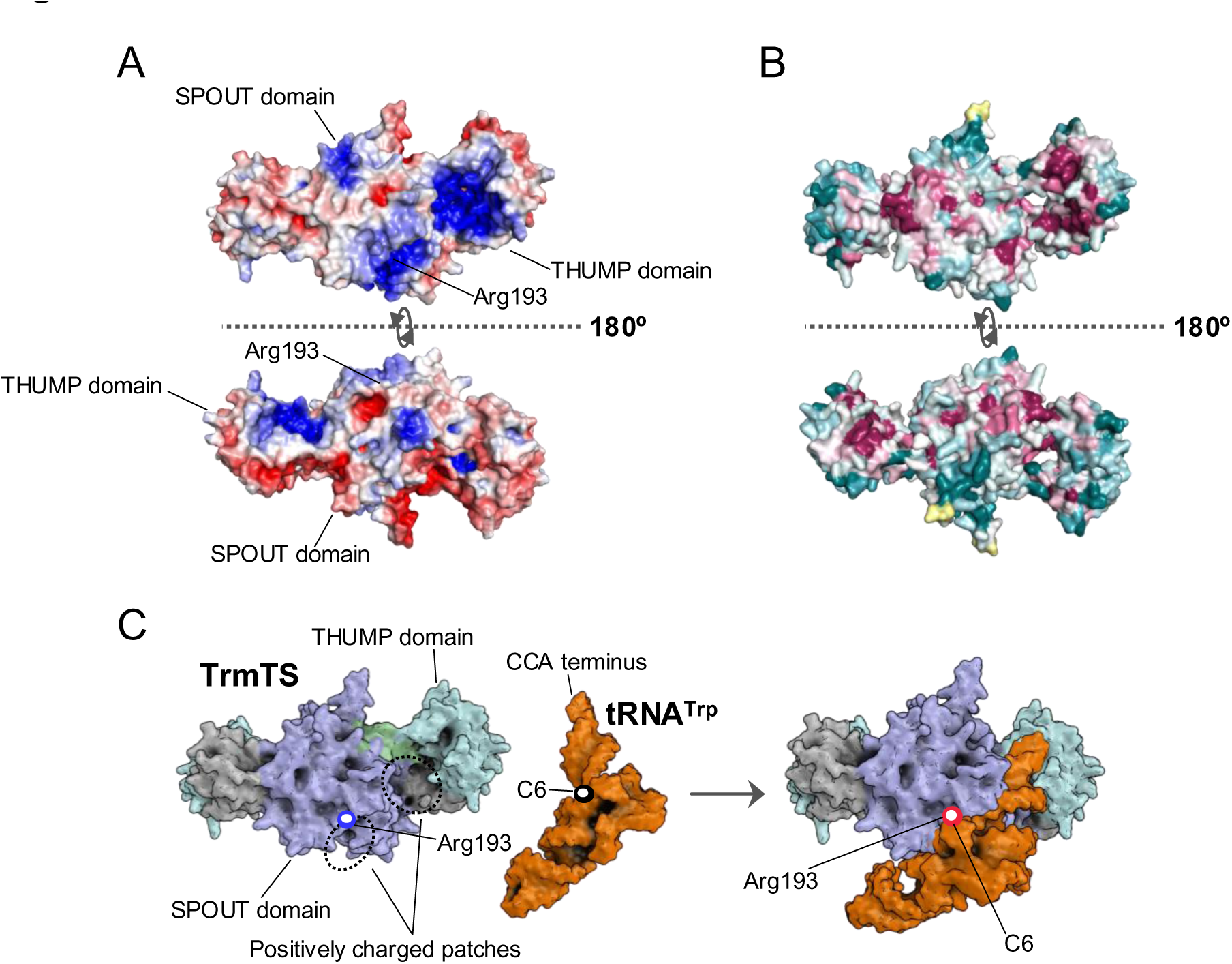
Hypothetical binding model between TrmTS and tRNA. A, Positive (blue) and negative (red) charged areas are mapped onto the surface of TrmTS dimer structure where +/– 5 kBT/e values were used for the visualization of the electrostatic potential map. B, Conserved and non-conserved amino acid regions are colored in magenta and dark green, respectively. C, A binding model between TrmTS and tRNA^Trp^ was predicted using AlphaFold3. The THUMP domain of TrmTS captures the CCA terminus of substrate tRNA and then the positively charged area of the SPOUT domain interact with the three-dimensional core of tRNA. Finally, the modification site (ribose of C6) in tRNA is introduced into the catalytic pocket of SPOUT domain and accesses the catalytic Arg193 residue.

## DISCUSSION

In this study, we have discovered a new tRNA methyltransferase using comparative genomics. This tRNA methyltransferase, TrmTS possesses highly unusual composition of domains, namely N-terminal ferredoxin-like, THUMP and SPOUT catalytic domains. Until this study, all THUMP-related tRNA methyltransferases reported possessed a Rossmann fold catalytic domain and nucleosides they produced were restricted to m^2^G and m^2^_2_G (56). Thus, our findings extend the concept of architecture of tRNA methyltransferases.

Although simple amino acid sequence alignment could not find the three conserved motifs in TrmH (Supplementary Fig. 5A), our mutagenesis study based on the structural comparison between the predicted TrmTS structure and TrmH identified the conserved amino acid residues which are important for the catalysis and structure. These results implies that the catalytic mechanism of TrmTS is probably common with other SPOUT RNA 2′-*O*-methyltransferases. However, one important residue is missing in TrmTS. The asparagine residue in the motif 1 of TrmH (Asn35 in *T. thermophilus* TrmH) is involved in the release of product (S-adenosyl-L-homocysteine) (26). However, the corresponding amino acid residue is not found in the sequence of TrmTS. Because Trm56 does not possess the asparagine residue as well (27), these archaeal enzymes (TrmTS and Trm56) may possess a different mechanism for the release of products. To clarify the details of roles of amino acid residues in TrmTS, a crystal structural study of the substrate-bound form of TrmTS is necessary. We found a TrmTS-specific linker region between the THUMP and SPOUT catalytic domains. Two lysine residues (Lys155 and Lys160) are highly conserved in the linker region of the TrmTS-like proteins. Lys160 in one subunit can interact with Asp270 residue in another subunit. Our mutagenesis study suggests that the Lys160 residue is important for the structure of TrmTS because the TrmTS Lys160Ala mutant protein was heavily degraded in *E. coli* cells.

Furthermore, the substitution of Lys155 with alanine decreases the Vmax value. Because the Lys155 residue is located far from the catalytic pocket, Lys155 residue may be involved in the movement of domains during the methylation process. Taking these results together, although the linker region between the THUMP and SPOUT domain is TrmTS-specific, this linker is essential for the activity of TrmTS.

TrmTS can form Am6 and Um6 as well as Cm6 in tRNA. Thus, the guanine base at position 6 is a negative determinant for the methylation by TrmTS. This enzymatic property is unique in tRNA 2′-*O*-methyltransferases. For example, TrmH (36), Trm56 (114) and archaeal TrmJ (37) strictly recognize G18, C56 and C32, respectively, in tRNA. Furthermore, TrmL recognizes only pyrimidine nucleosides at position 34 (115). Moreover, bacterial TrmJ does not recognize the base at position 32 (38). In the case of TrmTS, the 2-amino group of guanine base at position 6 in tRNA may cause steric hindrance around the catalytic pocket.

The THUMP-domain captures the CCA terminus. This concept has been established by numerous biochemical and structural studies (60, 63, 66, 75–81). In this study, we demonstrate that TrmTS, a novel THUMP-related tRNA methyltransferase, also requires the CCA terminal region for methylation. Furthermore, the deletion of the CCA terminal region from tRNA^Trp^ transcript causes the loss of affinity for TrmTS: the deletion mutant tRNA^Trp^ transcript does not work as a competitive inhibitor. Thus, our biochemical experiment shows that the interaction between the CCA terminus in tRNA and the THUMP domain of TrmTS is essential for the binding process. Our hypothetical model in Fig. 9C suggests that the three-dimensional core of substrate tRNA contacts the positively charged areas of TrmTS and then an induced-fit process occurs. To clarify the precise interaction between TrmTS and tRNA, the structural study of the complex between TrmTS and tRNA is necessary.

The physiological role of the Cm6 modification by TrmTS is unclear. Because TrmTS-like proteins are encoded in the genomes from thermophilic archaea and bacteria, we assumed that deletion of the *trmTS* gene might affect growth at high temperatures. However, the *ΔtrmTS* strain showed only a slight growth retardation at high temperatures. In *T. kodakarensis* tRNA^Trp^, numerous modified nucleosides exist (Fig. 1). Therefore, the Cm6 modification may work with other modifications at high temperatures. This idea is in line with the previous report that initiator tRNA^Met^ from *Pyrodictium occultum*, a hyper-thermophilic archaeon, is stabilized by 2′-*O*-methylation at multiple positions (41). Furthermore, in the case of *Pseudomonas aeruginosa*, the Am32 modification by TrmJ confers oxidative-stress resistance (42). Therefore, there is a possibility that the Cm6 modification by TrmTS may function in a stressful environment. To clarify the physiological role of Cm6 modification, further study is required.

## ACKNOWLEDGEMENTS

We thank the RI room staffs in the Advanced Research Support Center (Ehime University) for use of radioisotope compounds. Mass spectrometry experiments were conducted with the support of University of the Ryukyus, Research Facility Center.

## DATA AVAILABILITY

All data used in this study are available from the corresponding authors upon request.

## DATA

Supporting information is avairable at NAR Online: Supplementary Figures 1-8, and Supplementary Tables 1-6.

## CONFLICT OF INTERST

The authors declare no conflict of interest.

