## Supporting information for "A transfer RNA methyltransferase with an unusual domain composition catalyzes 2′-*O*-methylation at position 6 in tRNA"

**Transfer RNA methyltransferase that possesses an unprecedented domain composition catalyzes the 2’-*O*-methylation at position 6 in tRNA.**

**Teppei Matsuda^1^, Ryota Yamagami^1^*, Aoi Ihara^2^, Takeo Suzuki^3^, Akira Hirata^2^ and Hiroyuki Hori^1^***

1. Department of Materials Science and Biotechnology, Graduate School of Science and Engineering, Ehime University, 3 Bunkyo-cho, Matsuyama, Ehime 790-8577, Japan
2. Department of Natural Science, Graduate School of Technology, Industrial and Social Science, Tokushima University, 2-1 Minamijosanjimacho, Tokushima, Tokushima 770-8506, Japan
3. Department of Medical Biochemistry, Graduate School of Medicine, University of the Ryukyus, 207 Uehara, Nishihara, Okinawa 903-0125, Japan.

* To whom correspondence should be addressed:

Hiroyuki Hori

Department of Materials Science and Biotechnology, Graduate School of Science and Engineering, Ehime University, 3 Bunkyo-cho, Matsuyama, Ehime 790-8577, Japan.

Ryota Yamagami

Department of Materials Science and Biotechnology, Graduate School of Science and Engineering, Ehime University, 3 Bunkyo-cho, Matsuyama, Ehime 790-8577, Japan.

**MATERIALS AND METHODS**

**Optimization of KCl concentration for the methyl-transfer reaction by Tk1257 gene product**

Total RNA of *Escherichia coli* DH5α strain was prepared by phenol-chloroform extraction. Transfer RNA mixture was prepared from the total RNA by 10% PAGE (7 M urea). 0.5 μg Tk1257 gene product, 0.5 A260units *E. coli* tRNA mixture and 200 mM ^3^H-labeled SAM in 150 μL of buffer A [50 mM Tris-HCl (pH 7.6), 5 mM MgCl_2_, 6 mM 2-mrecaptoethanol and various concentrations of KCl] were incubated at 75°C. At 0, 5, 10, 30, 60 and 120 min-period, 25 μL of the reaction mixture was spotted onto a Whatman 3 MM filter. Incorporations of ^3^H-methyl group into *E. coli* tRNA mixture were assessed as described in the Materials and Methods section of the main text.

**Expression of TrmTS in the *ΔtrmTS* strain**

The Tk1765 (*chiA*) region in the *ΔtrmTS* strain was replaced by the *trmTS* gene (Tk1257) as reported previously (1).

**FIGURE LEGENDS**

**Supplementary Fig. 1.** ^3^H-methyl group incorporation velocities into *E. coli* tRNA mixture by Tk1257 gene product in the presence of 0, 100, 200, 300, 400 and 500 mM KCl are compared. The velocity in the presence of 100 mM KCl is expressed as 100%. The assay was independently replicated three times (n = 3).

**Supplementary Fig. 2.** Transfer RNA^Trp^ transcript was methylated by TrmTS and non-radioisotope labeled SAM and digested with RNase A. The resultant RNA fragments were analyzed by LC/MS. Before the methylation (left), 5’-HO-GGGGGCmGUp-3’ (m/z 876.79, z = -3) was not observed. In contrast, after the methylation (right), this fragment appeared.

**Supplementary Fig. 3.** **A**, Secondary structure of tRNA^Pro^ transcript is depicted in a cloverleaf structure. **B**, Transfer RNA^Pro^ transcript was methylated by TrmTS and non-radioisotope labeled SAM and digested with RNase T1. The resultant RNA fragments were analyzed by LC/MS. Before the methylation (left), 5’-HO-CCmGp-3’ (m/z 986.13, Z = -1) was not observed. In contrast, after the methylation (right), this fragment appeared.

**Supplementary Fig. 4.** Construction of the *ΔtrmTS* strain. **A**, KUWA strain was used as a parent strain and homologous recombination with a plasmid vector, pUTS, in which the selectable marker gene (*pdaD*; Tk0149) was inserted between the TK1256 and TK1258 genes. The plasmid pUTS, is not maintained as a plasmid in *T. kodakarensis* cells due to the lack of a replication origin. The pairs of oligonucleotides (TS-outF and TS-outR; TS-AGF and TS-outR; sequences in Supplementary Table 3) used to amplify genomic DNA from pUTS-generated agmatine-independent transformants are indicated. **B**, Electrophoresis of DNA molecules amplified by PCR from *T. kodakarensis* KUWA (lanes 1, 3 and 5) and *ΔtrmTS* (lanes 2, 4 and 6) using primer pairs TS-outF and TS-outR (lanes 1 and 2), or TS-AGF and TS-outR (lanes 3 and 4) or TS-AGF and TS-AGR (lanes 5 and 6). The DNA molecules amplified using primers TS-outF and TS-outR from *T. kodakarensis* KUWA and *∆trmTS* were, as predicted, 3.0 and 3.3 kb, respectively. Furthermore, the DNA molecules amplified using primers TS-AGF and TS-outR or TS-AGF and TS-AGR from the *ΔtrmTS* strain were only detected in the sample from the *ΔtrmTS* strain, exhibiting the insertion of *pdaD* gene between the TK1256 and TK1258 genes. Lane M contained size standards. **C**, Schematic drawing of the replacement of TK1765 region by TK1257 gene region. TK1257 gene product is expressed by TK2164 promoter in the *ΔtrmTS* strain. **D**, the replacement of Tk1765 region by TK1257 gene region was checked by PCR. The nucleotide sequence of amplified DNA was checked. **E**, Expression of TrmTS in the *ΔtrmTS* strain was confirmed by western blotting analysis.

**Supplementary Fig. 5.** Multiple sequence alignment and structural comparison between TrmTS model and TrmH. **A,** Multiple sequence alignment of the primary sequence of SPOUT domains from *T. thermophilus* TrmH, *E. coli* TrmL, *E. coli* TrmJ, *P. horikoshii* Trm56, and *T. kodakarensis* TrmTS was created with ClustalW. The motifs found in TrmH are denoted. **B**, Monomeric structures of SPOUT catalytic domains of TrmTS model and TrmH are provided. Conserved amino acid residues are highlighted in red. **C**, The dimer structure of SPOUT catalytic domains of TrmTS and TrmH are compared. The conserved residues are highlighted by stick models. To show the subunit interaction, amino acid residues in one subunit are colored in red and those in another subunit are colored in purple.

**Supplementary Fig. 6.** The dimer interfaces of SPOUT RNA 2’-O-methyltransferase are analyzed by the calculation of hydrophobic areas and topology diagrams. **A**, The hydrophobic areas highlighted in orange are calculated. The top structures are the overall dimeric structures of SPOUT RNA 2’-O-methyltransferase. The middle ones are the overall monomeric structures of SPOUT RNA 2’-O-methyltransferase. **B**, Topology diagrams are provided.

**Supplementary Fig. 7.** The gel-filtration column chromatography revealed that the SPOUT domain of TrmTS forms dimeric structure. **A**, Full-length of TrmTS protein was analyzed by Superdex-200 gel-filtration system. The size marker is denoted by an arrow. Roman numbers indicate fraction numbers. **B**, Proteins in each fraction from the gel-filtration experiment shown in panel A was analyzed by 15% SDS-PAGE. The proteins were visualized with CBB. **C**, the SPOUT domain of TrmTS was analyzed by Superdex-200 gel-filtration system. The size marker is denoted by an arrow. Roman numbers indicate fraction numbers. **D**, Proteins in each fraction from the gel-filtration experiment shown in panel C was analyzed by 15% SDS-PAGE.

**Supplementary Fig. 8.** RNA folding analysis revealed that truncated tRNA transcripts (**A**; transcript 6, **B**; transcript 7, **C**; transcript 8) are predicted to be fold into not typical cloverleaf structure of tRNA. RNAfold was used for the structure prediction.
