## Supplementary figures and images for "A transfer RNA methyltransferase with an unusual domain composition catalyzes 2′-*O*-methylation at position 6 in tRNA"

### Supplementary Figure 1

#### Supplementary Fig. 1

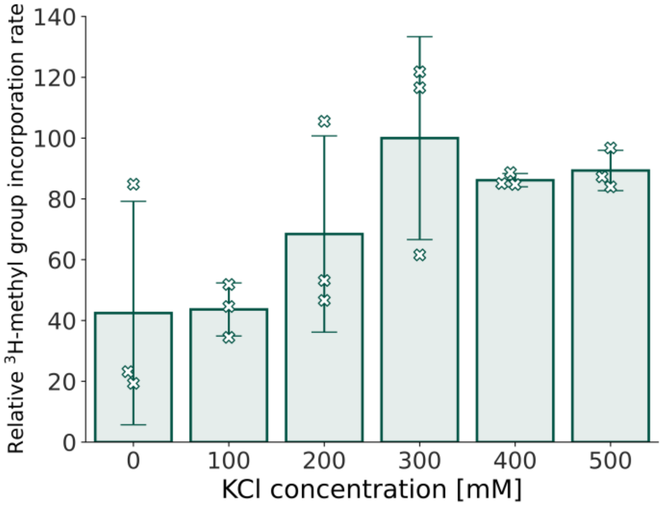

### Supplementary Figure 2

#### Supplementary Fig. 2

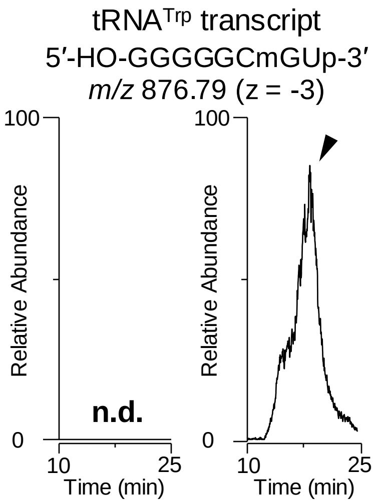

### Supplementary Figure 3

#### Supplementary Fig. 3

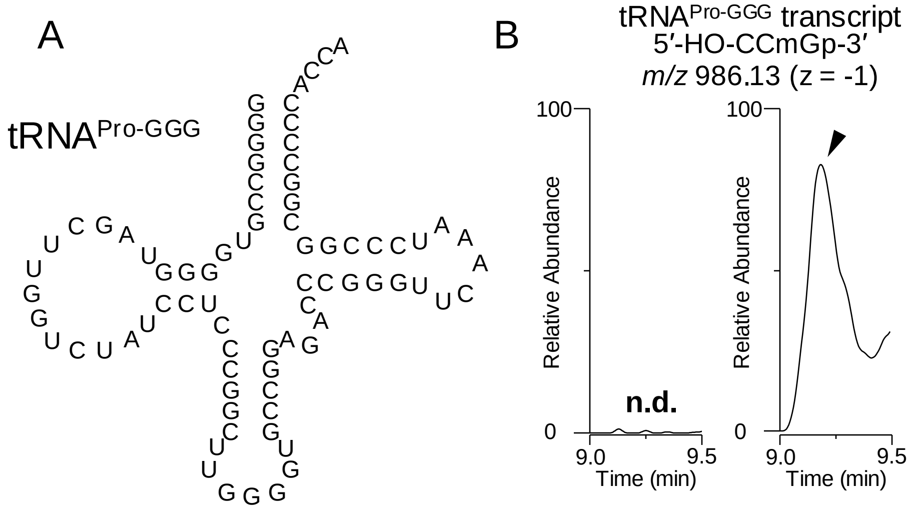

### Supplementary Figure 4

#### Supplementary Fig. 4

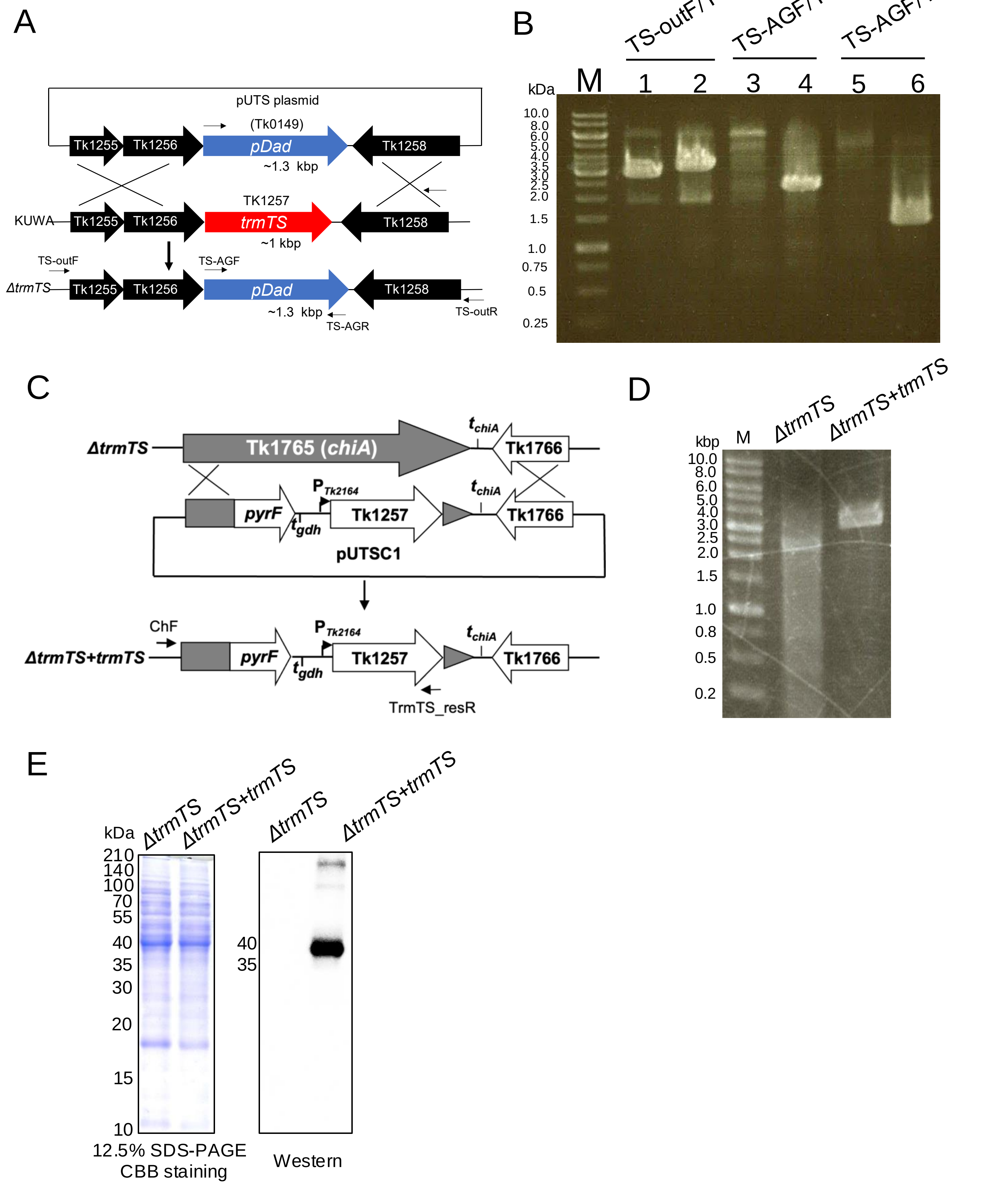

### Supplementary Figure 6

#### Supplementary Fig. 6

A

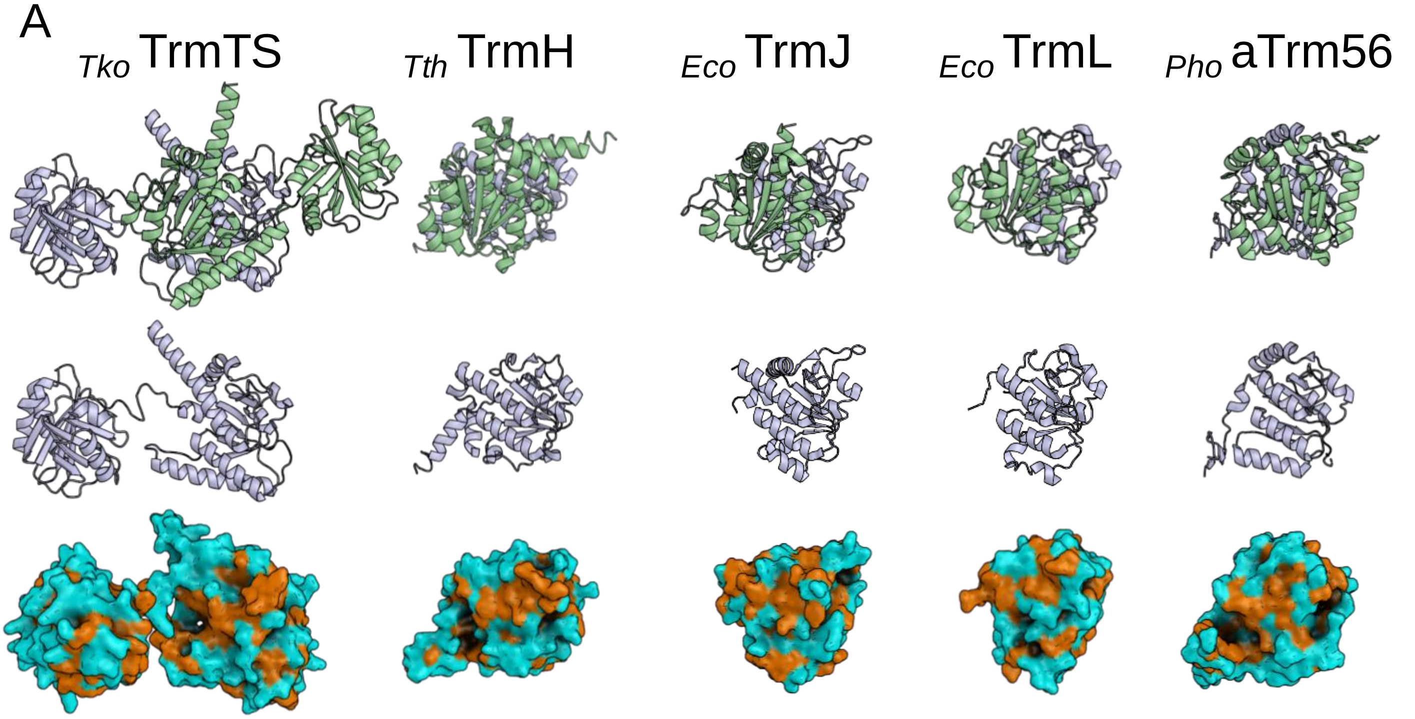


B

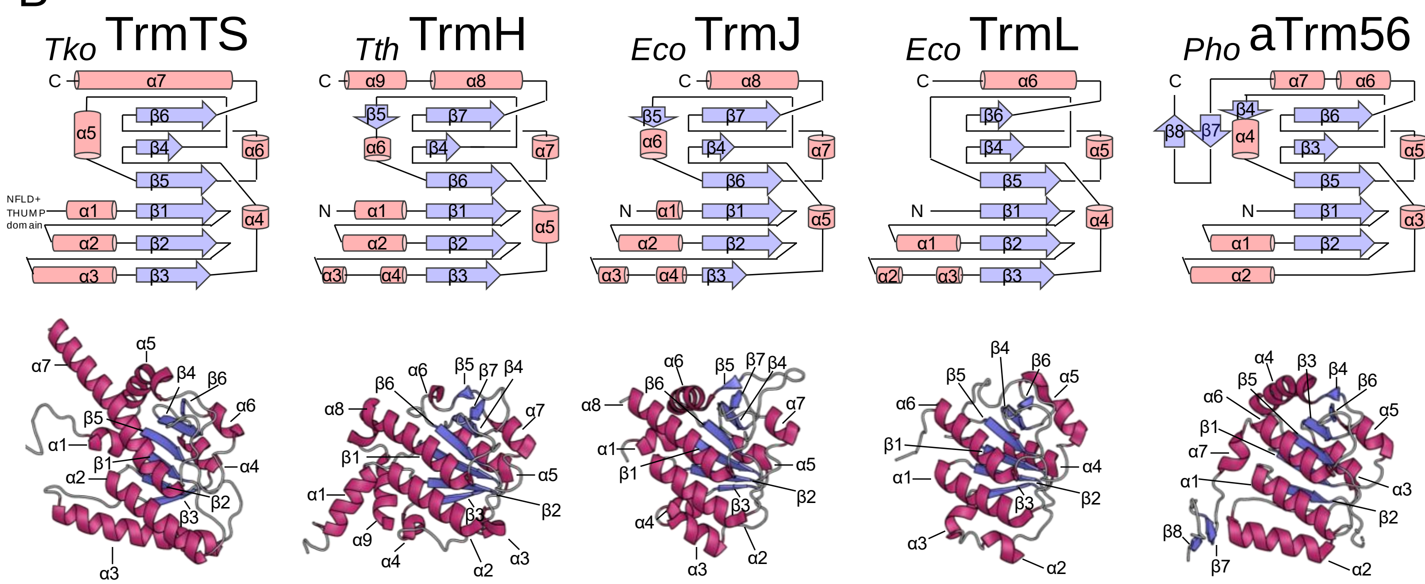

### Supplementary Figure 7

Supplementary Fig. 7

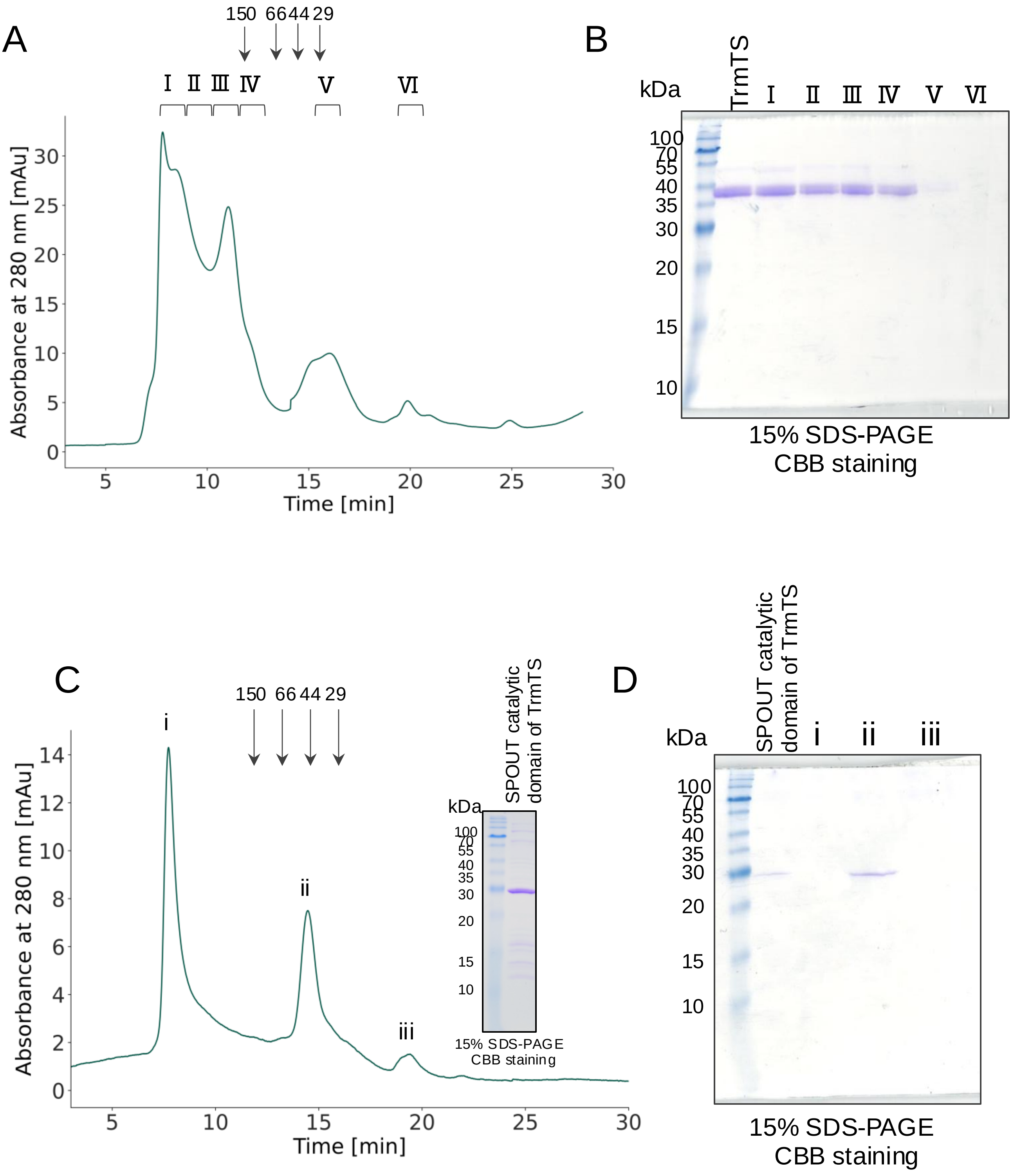

### Supplementary Figure 8

#### Supplementary Fig. 8

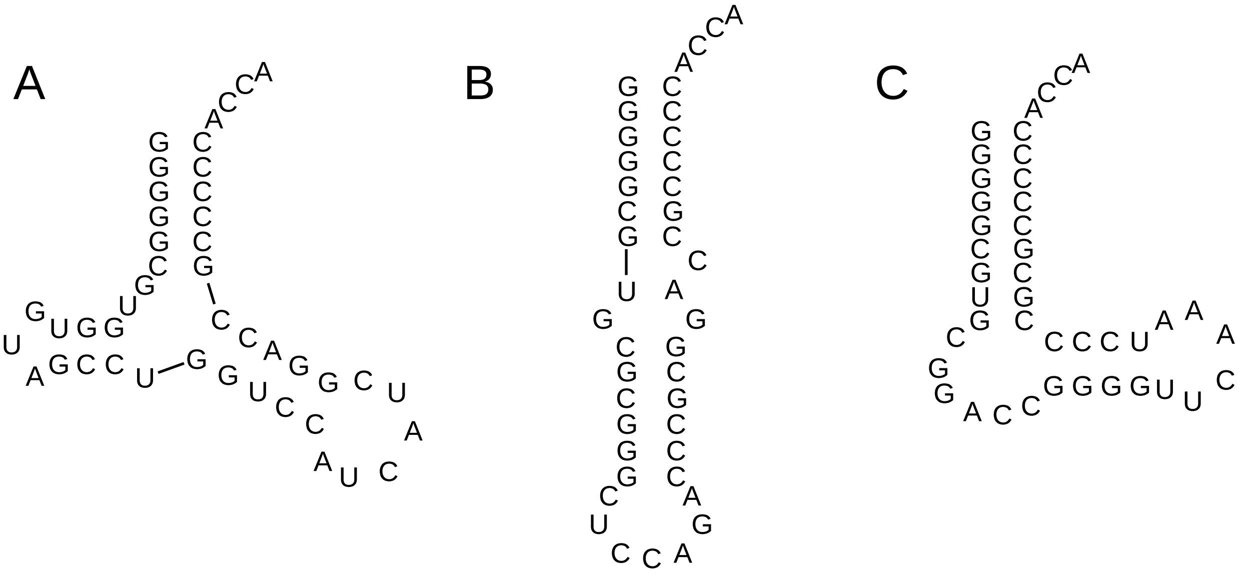


Transcript 6 | Transcript 7 | Transcript 8
