## Supplementary Figure 5 for "A transfer RNA methyltransferase with an unusual domain composition catalyzes 2′-*O*-methylation at position 6 in tRNA"

### Supplementary Fig. 5

A

```
                                                                    Motif 1
                  1        10        20        30        40        50
Tth_TrmH          MRERTEARRRRIEEVLRRRQPDLTVLLENVHKPHNLSAILRTCDAVGVLEAHAVNPTG..
Eco_TrmL          .....................MLNIVLYEPEIPPNTGNIIRLCANTG.FRLHIIEPMG..
Eco_TrmJ          ..................MLQNIRIVLVETSHTGNMGSVARAMKTMGLTNLWLVNPLVKP
Pho_Trm56         ....MIVVLRLGHRPERDKRVTTHVALTARAFGADGIIIASEEDEKVKESVEDVVKRWG.
Tko_TrmTS_SPOUT   ......EYRKYTPDKFDARKLFSKVTLVQMPYWGDYKACRKFGEKIGRAAQAFEVKELII

                          60        70        80        90        100
Tth_TrmH          ......GVPTFNETSG...GSHKWVYLRVHPDLHEAFRFLKERGFTVYATALREDARDFR
Eco_TrmL          ......FAWDDKRLRR...AGLDYHEFTAVTRHHDYRAFLEAENPQRLFALTTKGTPAHS
Eco_TrmJ          DSQAIALAAGASDVIG...NAHIVDTLDEALAGCSLVVGTSARSRTLPWPMLDPRECGLK
Pho_Trm56         .......GPFFIEFNR...NWRKVMKEFTGVKVHLTMYGLHVDDVIEELKEKLKKGEDFM
Tko_TrmTS_SPOUT   APKEKMDAYELMEFLRGIKIGQESRYQIQREAYPWKVEKVPVSVWDLYQVVRDKRRNKRL

                                  Motif 2             Motif 3
                      110       120       130       140       150       160
Tth_TrmH          EVDYTK..PTAVLFGAEKWGVSEEALALA..DGAIKIPMLGMVQSLNVSVAAAVILFEAQ
Eco_TrmL          AVSYQD..GDYLMFGPETRGLPASILDALPAEQKIRIPMVPDSRSMNLSNAVSVVVYEAW
Eco_TrmJ          SVAEAANTPVALVFGRERVGLTNEELQKC..HYHVAIAANPEYSSLNLAMAVQVIAYEVR
Pho_Trm56         IIVGAEKVPREVYELADYNVAIGNQPHSEVAALAVLLDRLLEGKGLKKEFKGAKIKIVPQ
Tko_TrmTS_SPOUT   LIITDPKGPTLAEVKEKLAKDMHYAKEVVIFIGSREGIPRGLFRFADYVVDLAPYMTFAT

                        170       180       190
Tth_TrmH          RQRLKAGLYDRPRLDPELYQKVLADWLRK...............................
Eco_TrmL          RQLGYPGAVLRD................................................
Eco_TrmJ          MAWLATQENGEQVEHEETPYPLVDDLERFYGHLEQTLLATGFIRENHPGQVMNKLRRLFT
Pho_Trm56         ARGKKVVEVQGYAEQDKAEGKATPGKNWEN.........SGFTGDNP.............
Tko_TrmTS_SPOUT   EHGIPATLVSLWEVYEEFLRKSEEKAEDSS..............................

Tth_TrmH          ............................
Eco_TrmL          ............................
Eco_TrmJ          RARPESQELNILRGILASIEQQNKGNKAE
Pho_Trm56         ............................
Tko_TrmTS_SPOUT   ............................
```

B

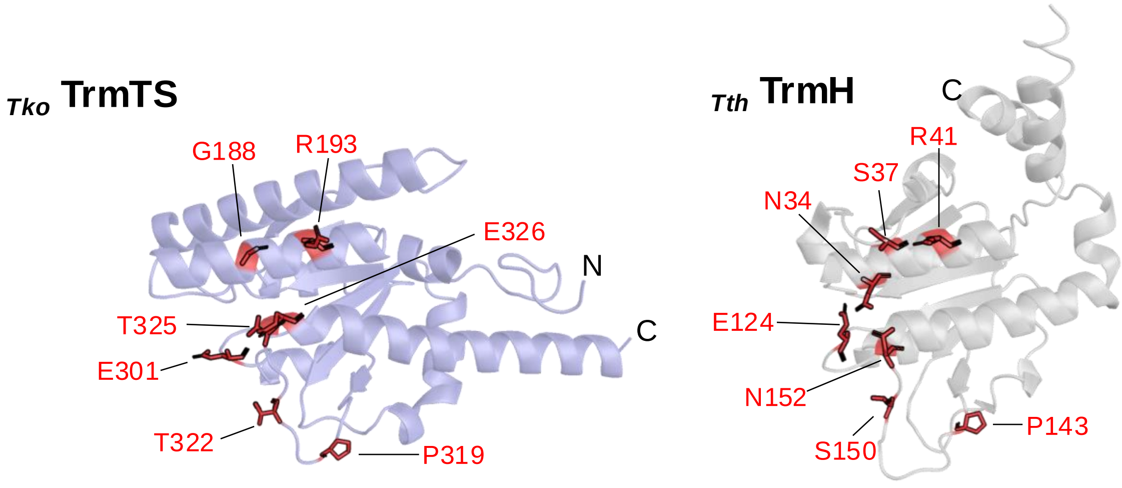


C

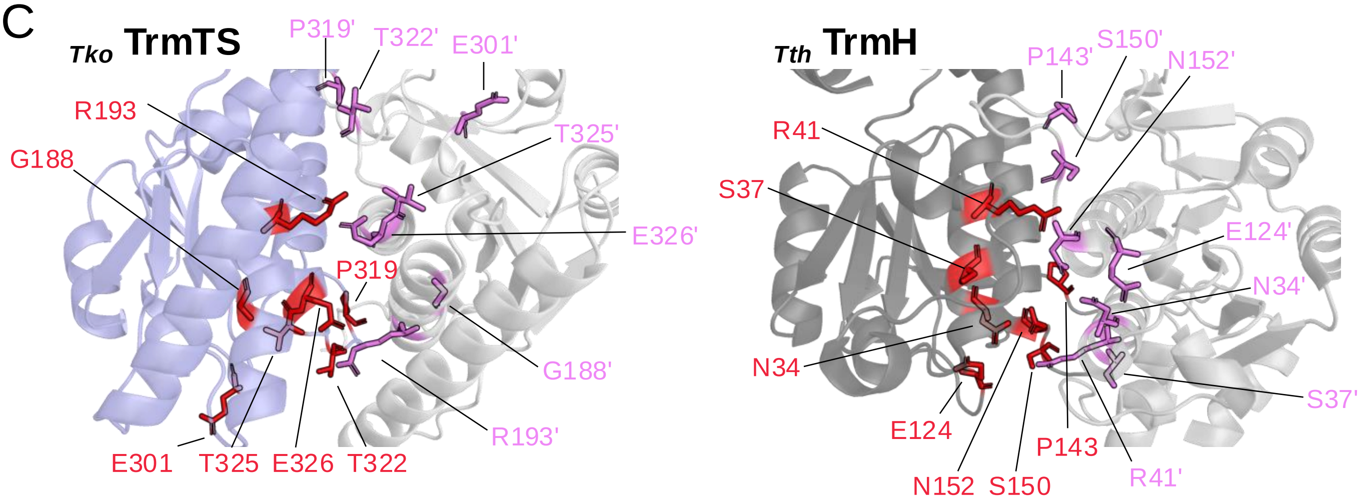
